# Parallel Distributed Networks Dissociate Episodic and Social Functions Within the Individual

**DOI:** 10.1101/733048

**Authors:** Lauren M. DiNicola, Rodrigo M. Braga, Randy L. Buckner

## Abstract

Association cortex is organized into large-scale distributed networks. One such network, the default network (DN), is linked to diverse forms of internal mentation, opening debate about whether shared anatomy supports multiple forms of cognition. Alternatively, subtle distinctions in cortical organization could remain to be resolved. Using within-individual analysis procedures that preserve idiosyncratic details of cortical anatomy, we probed whether multiple tasks from two domains - Episodic Projection and Theory of Mind (ToM) - rely upon the same or distinct networks. In an initial experiment (*n*=6, subjects scanned 4 times each), we found evidence that Episodic Projection and ToM tasks activate distinct functional regions distributed throughout cortex, with adjacent regions in parietal, temporal, prefrontal and midline zones. These distinctions were predicted by the hypothesis that the DN comprises two parallel, interdigitated networks. One network, linked to parahippocampal cortex (PHC), is preferentially recruited during Episodic Projection, including both remembering the past and imagining the future. A second juxtaposed network, which includes the temporoparietal junction (TPJ), is differentially engaged during multiple forms of ToM tasks. The TPJ-linked network is interwoven with the PHC-linked network in multiple zones, including the posterior and anterior midline, making clear why it is difficult to fully resolve the two networks in group-averaged or lower-resolution data. We replicated all aspects of this network dissociation in a second, prospectively acquired dataset (*n*=6). These results refine our understanding of the functional-anatomical organization of association cortex as well as raise questions about how functional specialization might arise in parallel, juxtaposed association networks.

## Introduction

Primate association cortex comprises multiple large-scale networks that each involve widely-distributed regions. Evidence for these networks comes from anatomical studies in non-human primates (e.g., Goldman-Rakic 1988; Mesulam 1990) and human functional neuroimaging studies that have broadly surveyed cortical network architecture (e.g., Doucet et al. 2011; Power et al. 2011; Yeo et al. 2011). The degree to which the networks are specialized and how they support distinct functions remain topics of active investigation. One debate surrounds the role that the distributed network at the transmodal apex of association cortex^1^, often referred to as the default network (DN), plays in diverse forms of internally-generated mentation and cognitive processes more broadly. The DN is situated within zones of association cortex that are disproportionately expanded in hominin evolution and late to mature during development, making them particularly intriguing targets for understanding how distributed cortical networks come to possess specialization important to advanced aspects of human cognition (Buckner and Krienen 2013; Hill et al. 2010).

The canonical DN includes regions in the medial and lateral prefrontal cortex (PFC), lateral temporal cortex, posterior cingulate / retrosplenial cortex (PCC/RSC), medial temporal lobe (MTL) and inferior parietal lobule (IPL; e.g., Buckner et al. 2008; Gusnard and Raichle 2001). Tasks exhibiting activation that overlaps with the DN include autobiographical memory (Svoboda et al. 2007), future prospection (Schacter et al. 2012), social inference (Iacoboni et al. 2004; Mars et al. 2012; Schurz et al. 2014), and self-referential processing (e.g., D’Argembeau et al. 2005; Gusnard and Raichle 2001). Several early examinations of DN function highlighted convergence and shared patterns of activation across these multiple task domains. Commonalities were noted, for example, in the regions recruited for tasks targeting episodic memory, future prospection and representation of others’ mental states, also termed theory of mind (ToM; Buckner and Carroll 2007; Buckner et al. 2008; Spreng et al. 2009; see also Schacter et al. 2007).

Subsequent group studies incorporating within-subject task manipulations found differential recruitment of certain DN regions by distinct tasks (Andrews-Hanna et al. 2010; Andrews-Hanna et al. 2014a; DuPre et al. 2016; Rabin et al. 2010; Spreng and Grady 2010; see also Rosenbaum et al. 2007). In particular, retrieving episodic memories recruits regions near RSC and parahippocampal cortex (PHC); in contrast, inferring others’ mental states recruits a region of the IPL, extending into the temporoparietal junction (TPJ), that is rostral to the caudal IPL zone most typically associated with episodic remembering (Andrews-Hanna et al. 2014a; DuPre et al. 2016; Rabin et al. 2010). In a resultant organizational hypothesis of the DN, two subsystems were proposed featuring distinct regions, including dorsomedial PFC and MTL, as well as shared – or core – regions along the anterior and posterior midline (Andrews-Hanna et al. 2010; Andrews-Hanna et al. 2014b; see also Hassabis and Maguire 2007).

Recent technological advances and procedures for sampling individuals have allowed for better appreciation of the detailed spatial components of distributed association networks (Braga and Buckner 2017; Braga et al. 2019; Fedorenko et al. 2012; Gordon et al. 2017; Huth et al. 2016; Kong et al. 2018; Laumann et al. 2015; Michalka et al. 2015). Relevant to the study of the DN, Braga and Buckner (2017) revealed that two closely interdigitated networks are distributed within the bounds of the canonical DN (see also Braga et al. 2019; Buckner and DiNicola In Press). These dissociable networks, termed Network A and Network B for convenience, are interwoven but distinct across multiple zones of cortex. Certain spatial distinctions are consistent with those previously highlighted at the group level, such as the separation of rostral and caudal regions of the IPL, previously linked differentially to mentalizing and mnemonic functions (e.g., Andrews-Hanna et al. 2014a). However, individualized analyses also suggest new features that may be critical to functional understanding. First, the network distinctions are not restricted to local zones, but rather have dissociable components throughout the distributed extent of the networks. Second, the anatomical dissociations within individuals include regions along the anterior and posterior midline that were previously proposed as part of the DN “core” due to their participation in multiple task domains. The network organization as revealed within the individual, as well as findings from task-based activation patterns along the posterior midline within individuals (Peer et al. 2015; Silson et al. 2019), suggest that these zones may possess spatially separate functional regions. Altogether, these results raise the possibility that there may be broad functional specialization of the widely distributed networks, rather than local subzones of specialization that converge on core hubs.

Motivated by these possibilities and by the general goal of increasing our understanding of functional organization, we explored specialization of networks located within the bounds of the canonical DN using repeated scanning and approaches focused on characterizing idiosyncratic details within individuals. We first conducted intrinsic functional connectivity analysis to identify Networks A and B within each individual. We then examined task response patterns during multiple task contrasts targeting either Episodic Projection (i.e., remembering the past and imagining the future) or ToM (also called mentalizing) in order to explore functional-anatomical specialization of Networks A and B within the individual. Two independent samples were collected, each including 6 intensively sampled participants. Analysis techniques were finalized during analysis of the first sample (Exp. 1) and then applied during analysis of the second sample (Exp. 2), which was collected as a fully independent, prospective replication.

To foreshadow the results, Networks A and B exhibit replicable functional dissociation across distributed cortical zones. PHC-linked Network A is preferentially recruited for tasks requiring Episodic Projection, while TPJ-linked Network B exhibits preferential recruitment for tasks requiring ToM. Dissociation extends to topographically separate regions within the anterior and posterior midline and throughout the distributed zones of association cortex, pointing toward a revised understanding of how distributed networks may be organized to support distinct task processing demands.

## Experiment 1 Methods: Initial Functional Dissociation

### Participants

6 healthy adults, aged 18-32 (*Mean* = 22.2 years (SD = 2.4), 2 males, 5 right-handed) were recruited from the Boston area. All participants were native English speakers and screened to exclude neurological or psychiatric illness. Each participant was scanned across 4 separate MRI sessions, designed to chart (1) individualized network organization through analysis of intrinsic functional connectivity and (2) task-based activation through tests of Episodic Projection and ToM. Participants provided written informed consent using procedures approved by the Institutional Review Board of Harvard University and were paid for participation.

### MRI Data Acquisition

Scanning was conducted at the Harvard Center for Brain Science using a 3T Siemens Prisma-fit MRI scanner and a 64-channel phased-array head-neck coil (Siemens Healthcare, Erlangen, Germany). Foam padding provided head comfort and immobilization. Each scanning session was conducted on a separate (non-consecutive) day and lasted up to 2 hours. Participants viewed rear-projected stimuli through a mirror attached to the head coil. Prior to each session, the viewing location was adjusted so that the central point of the participant’s field of view was at a comfortable angle.

During each session, a rapid T1-weighted structural image with 1.2mm isotropic voxels was acquired first, using a multi-echo magnetization prepared rapid acquisition gradient echo (ME-MPRAGE) sequence (TR=2200ms, TE=1.57, 3.39, 5.21, 7.03ms, TI=1100ms, 144 slices, flip angle=7°, matrix=192 × 192 × 144, in-plane GRAPPA acceleration 4). T2-weighted blood oxygenation level-dependent (BOLD) runs were then acquired using a multi-band gradient-echo echo-planar pulse sequence (see Setsompop et al. 2012), generously provided by the Center for Magnetic Resonance Research at the University of Minnesota (TR=1000ms, TE=32.6ms, flip-angle 55°, 2.4mm isotropic voxels, multi-slice 5× acceleration, matrix=88 × 88 × 65, 65 slices covering cerebral cortex and cerebellum). A dual-gradient-echo B_0_ field map (2.4mm isotropic resolution, TR=295ms, flip angle=64°, Phase: 65 frames, Magnitude: 130 frames, TE_1_=4.45ms, TE_2_=6.91ms) was also acquired during each session to allow for correction of susceptibility-related inhomogeneities. Participants’ eyes were monitored and video-recorded using the Eyelink 1000 Core Plus with Long-Range Mount (SR Research, Ottawa, Ontario, Canada), and alertness was scored during each BOLD run.

### Fixation Runs for Intrinsic Functional Connectivity Analysis

Eight 7m 2s BOLD fixation runs were acquired per individual (approximately 56m of data; 2 runs per session). During the fixation task, participants were instructed to stay alert and still, and maintain fixation on a centrally presented black plus sign on a light grey background. Fixation runs occurred intermixed with other task paradigms. Fixation data were used for functional connectivity analysis in order to estimate network organization independent from the data used in subsequent task analysis.

### Task Paradigms

Task contrasts were selected that preferentially isolated either Episodic Projection or ToM demands. These task domains, albeit complex within themselves, were hypothesized to differentially recruit parallel networks that fall within the bounds of the default network (e.g., Andrews-Hanna et al. 2010; Andrews-Hanna et al. 2014a; DuPre et al. 2016). Two variants of each task type were constructed based on prior literature, maintaining the task structure and contrast strategy of the original work. Two variants were used, rather than more trials of the same type, to aim for generality and also to increase the number of unique rich stimuli. Participants were given detailed instructions and practice trials for each task during an initial training session. Instructions were repeated prior to each scanning session.

The Episodic Projection tasks involved episodic remembering of the past (PAST SELF) and episodic prospection of the future (FUTURE SELF; Andrews-Hanna et al. 2010). These tasks were contrasted with a common control condition (PRESENT SELF). The ToM tasks involved representing others’ mental states, either through making inferences about another person’s beliefs (FALSE BELIEF) or considering another person’s emotional pain or suffering (EMO PAIN; Dodell-Feder et al. 2011; Jacoby et al. 2016; Saxe and Kanwisher 2003); each was contrasted to its own matched control condition (FALSE PHOTO or PHYS PAIN). Example stimuli are provided in Table 1.

**Table 1.**
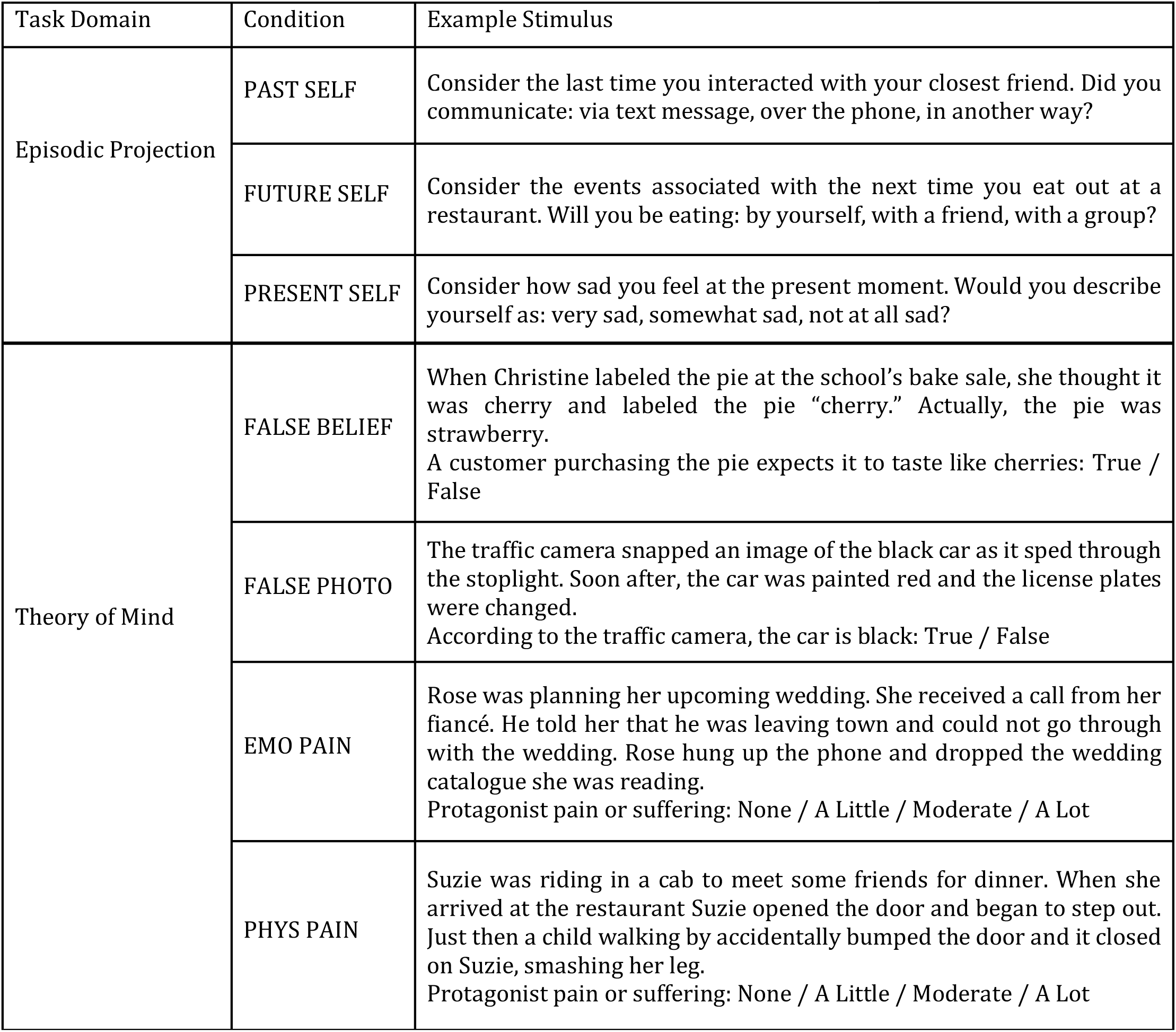
Sample stimuli from conditions of interest across primary tasks.

| Task Domain | Condition | Example Stimulus |
| --- | --- | --- |
| Episodic Projection | PAST SELF | Consider the last time you interacted with your closest friend. Did you communicate: via text message, over the phone, in another way? |
|  | FUTURE SELF | Consider the events associated with the next time you eat out at a restaurant. Will you be eating: by yourself, with a friend, with a group? |
|  | PRESENT SELF | Consider how sad you feel at the present moment. Would you describe yourself as: very sad, somewhat sad, not at all sad? |
| Theory of Mind | FALSE BELIEF | When Christine labeled the pie at the school's bake sale, she thought it was cherry and labeled the pie "cherry." Actually, the pie was strawberry.<br>A customer purchasing the pie expects it to taste like cherries: True / False |
|  | FALSE PHOTO | The traffic camera snapped an image of the black car as it sped through the stoplight. Soon after, the car was painted red and the license plates were changed.<br>According to the traffic camera, the car is black: True / False |
|  | EMO PAIN | Rose was planning her upcoming wedding. She received a call from her fiancé. He told her that he was leaving town and could not go through with the wedding. Rose hung up the phone and dropped the wedding catalogue she was reading.<br>Protagonist pain or suffering: None / A Little / Moderate / A Lot |
|  | PHYS PAIN | Suzie was riding in a cab to meet some friends for dinner. When she arrived at the restaurant Suzie opened the door and began to step out. Just then a child walking by accidentally bumped the door and it closed on Suzie, smashing her leg.<br>Protagonist pain or suffering: None / A Little / Moderate / A Lot |

#### Episodic Projection Tasks

Episodic Projection tasks and their control extended from a paradigm developed by Andrews-Hanna et al. (2010). The tasks involved visually presenting questions about hypothetical past or future scenarios (context-setting statements) and then providing three possible answer choices. Questions and answers targeted real-world experiences rather than unlikely or fantasy scenarios to maximize the likelihood that individuals would experience Episodic Projection.

The anchor for designing the Episodic Projection task contrasts was a prior observation that contrasting episodic future projection against a control of present self-reflection preferentially activated components of Network A (see Fig. 4 in Andrews-Hanna et al. 2010). Critically, both the target and control tasks involve self-referential processing, removing this dimension (to a degree) from the contrast. For the current study, we replicated the future-oriented condition used by Andrews-Hanna and colleagues (FUTURE SELF), and also created a new, parallel task condition focused on remembering the past (PAST SELF).

**Figure 1.**
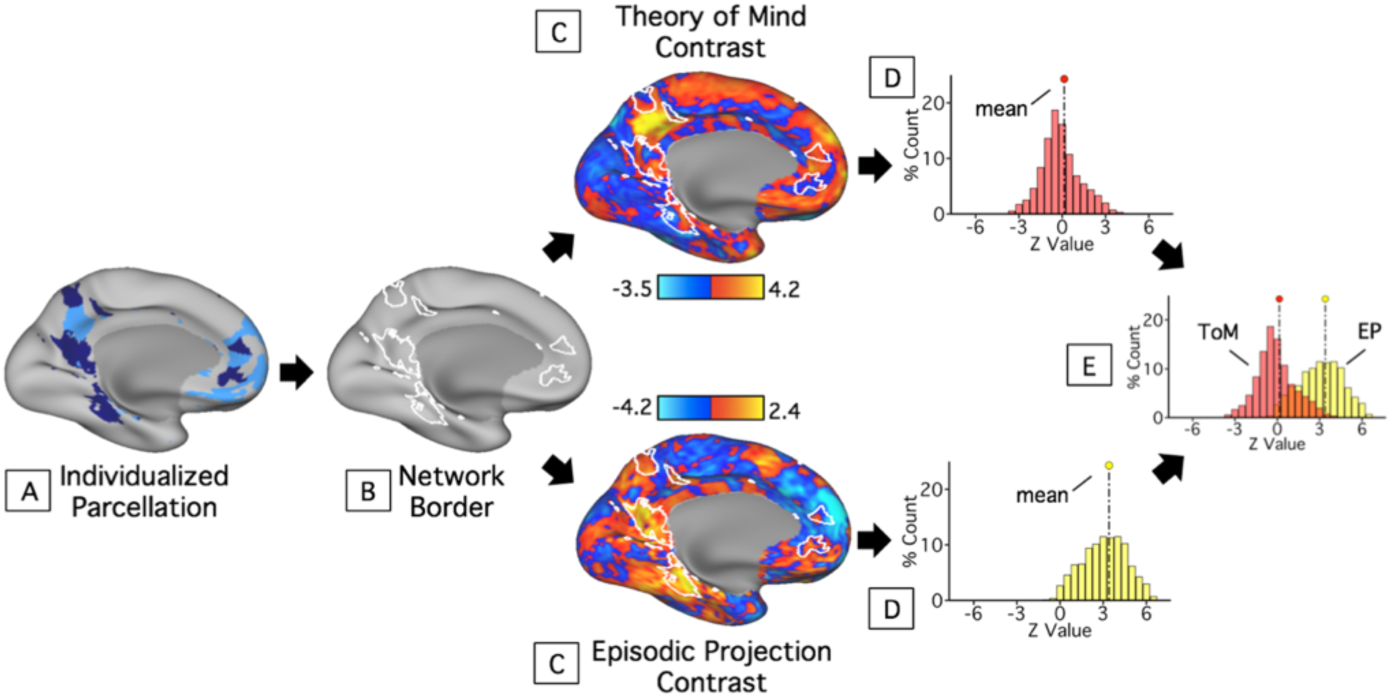
Procedure for testing functional dissociation within individuals. Within each subject, Networks A and B were identified using k-means clustering, as illustrated in (A). Each network’s border was defined (B) and overlaid on the unthresholded contrast maps for each task domain (C). The distribution of *z*-weighted values within each network’s boundaries was extracted and plotted (D). For each network, plots from both task domains were then visualized on a single graph (Theory of Mind, ToM, = red; Episodic Projection, EP, = yellow), and potential differences between the domain distribution means were quantified using effect sizes (E). While this figure illustrates distributions for Network A only, the procedure was performed identically for both Networks A and B.

**Figure 2.**
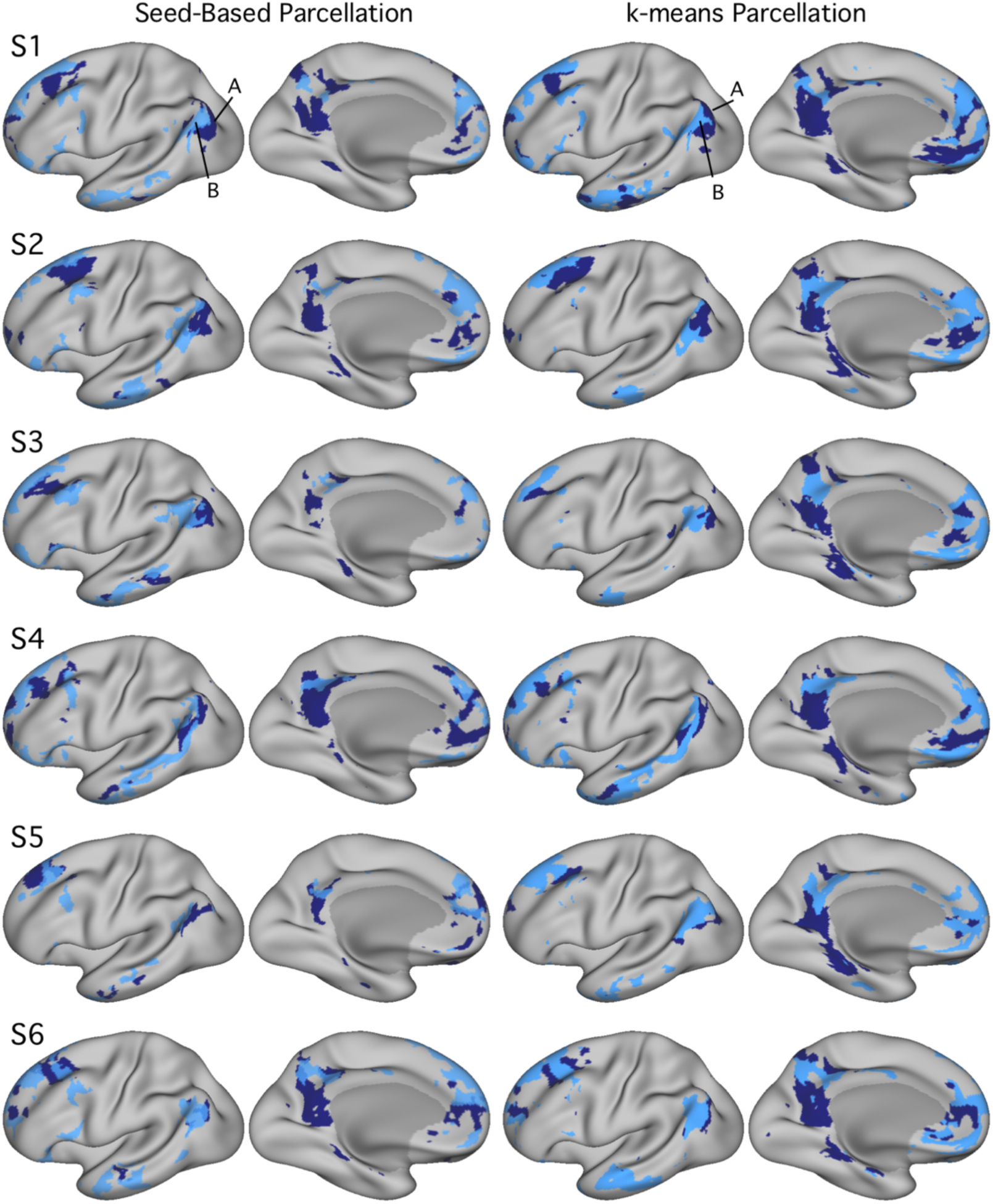
Networks A and B identified within individuals using both seed-based and k-means parcellation strategies. Estimates of Networks A and B from Exp. 1 identified using seed-based (left column) and k-means (right column) methods exhibit comparable maps. Although exact network boundaries differ by method, both techniques reveal interdigitated network patterns along the medial prefrontal and lateral frontal cortex, as well as similarly juxtaposed network boundaries along the posteromedial cortex and inferior parietal lobule in all individuals. *k*=17 for all k-means parcellations shown here; Network A appears in navy and Network B in light blue. Seed-based maps are thresholded at *r*=0.40.

**Figure 3.**
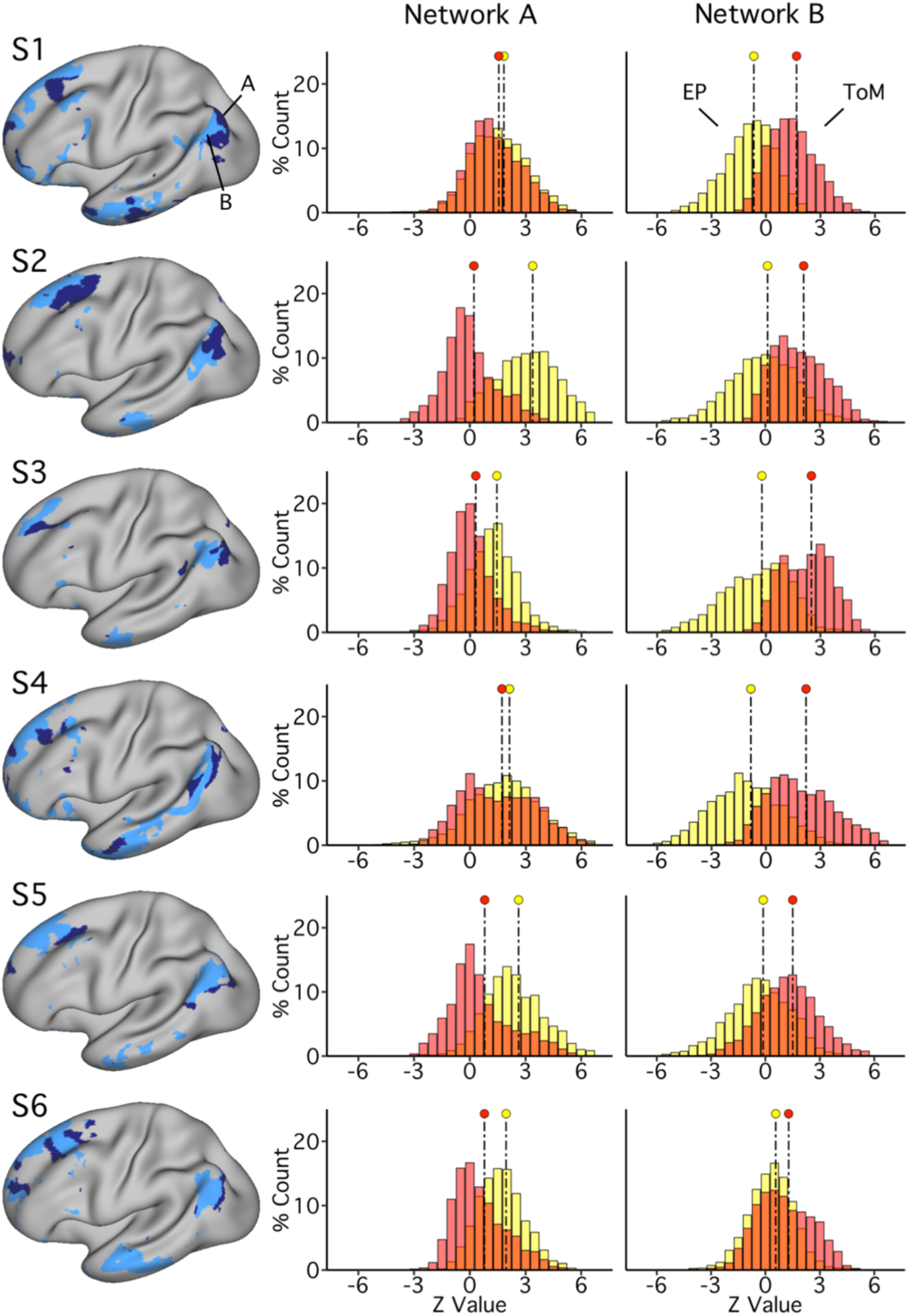
Networks A and B show functional dissociation within individuals in Exp. 1. The left column displays Network A (navy) and Network B (light blue) for each subject as defined by k-means clustering. The distributions in the right columns plot the functional responses within each network for the two task domains (ToM = red, Episodic Projection = yellow, Overlap = orange). See Fig 1 for method. For Network A, all 6 individuals reveal a functional response increase for Episodic Projection over ToM – and most strongly in 4 subjects (with Cohen’s *d* range = 0.88-2.03 for 4 subjects; *d=*0.19 for S1 and *d=*0.20 for S4). For Network B, all 6 individuals reveal the opposite pattern. The ToM response is increased over Episodic Projection (Cohen’s *d* range = 0.51-1.76). The consistent opposing patterns between networks are evidence for functional double dissociation.

**Figure 4.**
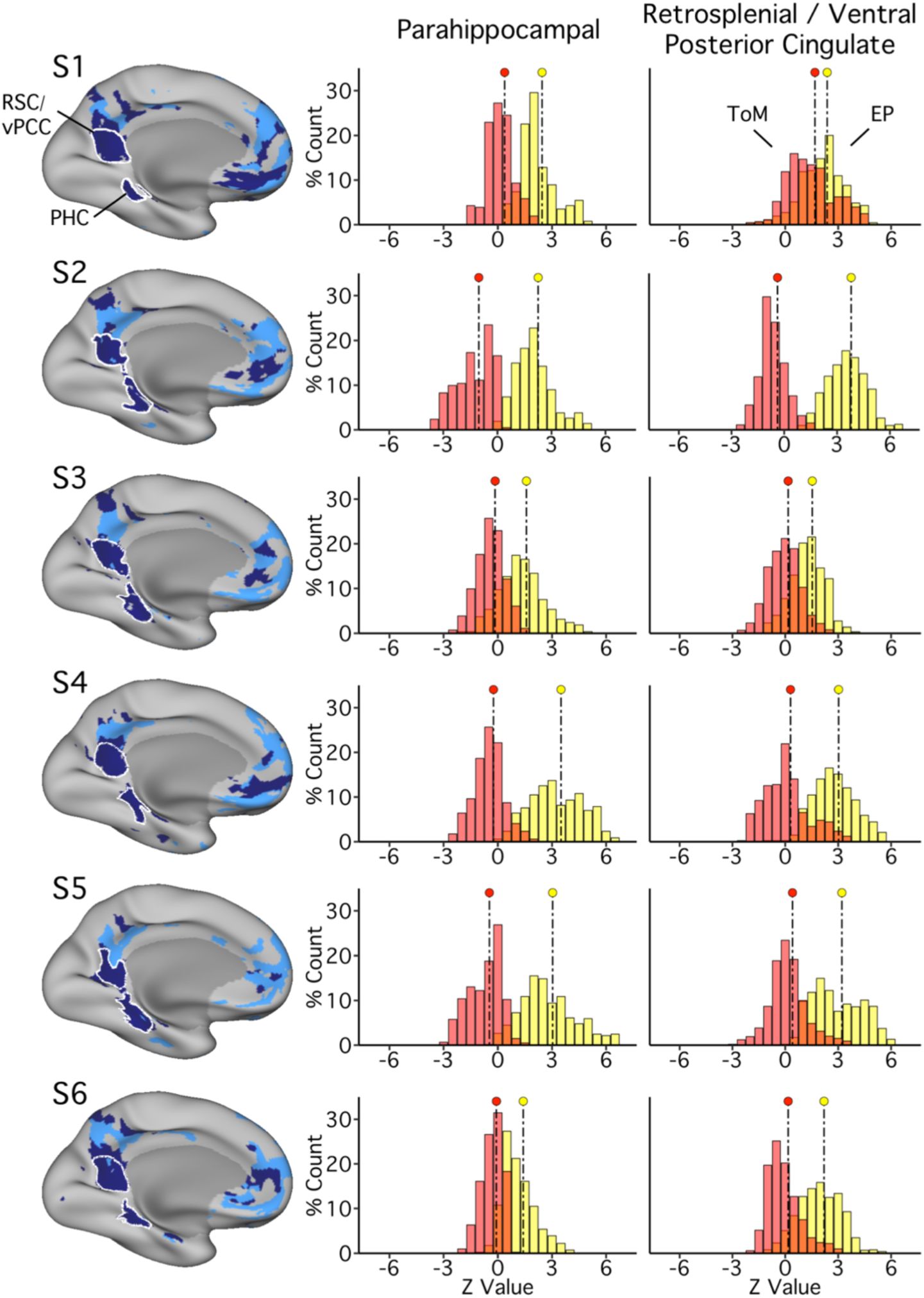
PHC and ventral PCC/RSC regions of Network A exhibit robust functional dissociation in Exp. 1. The left column illustrates parahippocampal (PHC) and retrosplenial / ventral posterior cingulate (RSC / vPCC) regions, as defined in each subject, outlined in white. The distributions plot the functional responses within each region of Network A for the two task domains (ToM = red, Episodic Projection = yellow, Overlap = orange). In all subjects for both regions, a clear functional response increase is evident for Episodic Projection over ToM (Cohen’s *d* ranges: RSC/vPCC: 0.54-4.43; PHC: 1.67-3.24).

The control task extended from the PRESENT SELF condition used by Andrews-Hanna et al. (2010), which primarily featured items directed at conceptual beliefs about oneself – issues in one’s life, feelings and beliefs – while other items concerned immediate perceptual experiences, such as physical feelings within the scanner. In the current experiment, statements about one’s conceptual beliefs, feelings and ideas were probed (PRESENT SELF). A separate condition then queried present physical and perceptual experiences (PRESENT PERCEPTION), and a final condition included parallel statements and questions about nonpersonal semantic knowledge (SEMANTIC). The task contrasts relevant to isolating regions suggestive of Network A were PAST SELF versus PRESENT SELF, and FUTURE SELF versus PRESENT SELF.

The full sets of task conditions were intermixed to create randomized trial orders that alternated across runs. Each 612s run included 30 trials (6 per condition). A run began with 12s of fixation for T1-stabilization. 30 trials were then presented sequentially. Each trial lasted 20s, with 5s of fixation, 10s during which the context-setting statements and answers were presented, and then 5s of additional fixation. Sentence structure, words and character length were matched across conditions. One run of 30 trials was collected during each MRI session, yielding 4 total runs (120 individual trials, 24 of each type; 40m of total Episodic Projection data).

#### Theory of Mind Tasks

ToM tasks and their control conditions extended from the False Belief and the Emotional/Physical Pain Stories paradigms (Dodell-Feder et al. 2011; Jacoby et al. 2016; Saxe and Kanwisher 2003), each of which targets representation of others’ mental states (e.g., Saxe 2006). These task contrasts were chosen because they show convergent patterns of activation that preferentially align with regions suggestive of Network B, in particular robust activation of the TPJ (Jacoby et al. 2016).

Each of the False Belief ToM runs included 10 trials, 5 featuring stories about characters with potentially false beliefs (FALSE BELIEF) and 5 control stories about objects (e.g., photographs or maps) with potentially false information (FALSE PHOTO; Dodell-Feder et al. 2011). The task contrast relevant to isolating Network B was FALSE BELIEF versus FALSE PHOTO. A run began with 12s of fixation for T1-stabilization. Each story was presented for 10s, followed by a true or false statement, presented for 5s. A 15s fixation period then occurred before the next trial began. Participants were asked to pay close attention to the details of each presented story and then respond to the statement. Stimuli included those from Dodell-Feder et al. (2011), as well as additional stimuli generously provided by the Saxe Laboratory.

Each of the Emotional/Physical Pain Stories runs (subsequently abbreviated “Other Pain”) included 10 trials, with 5 featuring stories containing an emotionally painful event (EMO PAIN) and 5 control stories containing a physically painful event (PHYS PAIN; Bruneau, Pluta and Saxe 2012; Jacoby et al. 2016). The task contrast relevant to isolating Network B was EMO PAIN versus PHYS PAIN. The timing of the runs and trials were identical to those described for the False Belief ToM runs. However, the structure of the questions and responses differed. During the question period participants selected, on a scale from 1-4, the amount of pain the protagonist of the story had experienced (numbers corresponded to ‘None’, ‘A Little’, ‘Moderate’ or ‘A Lot’). Stimuli were selected from the full set provided by Bruneau, Pluta and Saxe (2012). For each ToM task, a fixed randomized trial order alternated across sessions. One run of each ToM task paradigm was collected during each MRI session, yielding 8 total runs (4 per task, 40m of total ToM data).

### Exclusion Criteria and Quality Control

Each BOLD functional run was examined individually for quality. Scan and behavioral performance were assessed, and, for runs featuring skipped trials, eye videos were reviewed. Since the Episodic Projection and the Other Pain tasks involved subjective decisions, for which accuracy could not be calculated, behavioral performance metrics quantified (1) frequency of skipped trials and (2) average response times (RT; compared across conditions within each task).

Exclusion criteria included: (1) maximum absolute motion greater than 2mm, (2) signal to noise ratio less than or equal to 135, and (3) eyes closed during skipped task trials (see Table 2). Overall, 2 of 120 runs were excluded in Exp. 1 for motion (1 run for each of 2 subjects). No runs were excluded based on behavioral performance metrics (see Exp. 1 Results and Table 3 for details). Data quality was excellent, with mean maximum motion displacement less than 1mm for each subject.

**Table 2.** Quality control metrics.

|  | Subject | Maximum Motion (mm) | Signal-to-Noise Ratio | Excluded Runs |
| --- | --- | --- | --- | --- |
| Exp. 1 | S1 | 0.74 (0.33) | 256.75 (48.49) | 0 |
|  | S2 | 0.34 (0.14) | 316.95 (26.74) | 0 |
|  | S3 | 0.36 (0.20) | 305.90 (48.37) | 0 |
|  | S4 | 0.67 (0.23) | 203.20 (32.28) | 0 |
|  | S5 | 0.99 (0.46) | 226.00 (45.06) | 1 [ToM: motion] |
|  | S6 | 0.55 (0.34) | 248.79 (35.77) | 1 [EP: motion] |
| Exp. 2 | S7 | 0.41 (0.31) | 264.23 (50.40) | 0 |
|  | S8 | 0.36 (0.14) | 252.64 (28.99) | 0 |
|  | S9 | 0.71 (0.34) | 285.97 (44.09) | 0 |
|  | S10 | 0.56 (0.25) | 248.97 (30.34) | 3 [FIX: motion] |
|  | S11 | 0.74 (0.28) | 254.91 (34.84) | 2 [1 FIX: motion; 1 ToM: pres. error] |
|  | S12 | 0.30 (0.07) | 301.70 (42.14) | 1 [ToM: pres. error] |

Notes: Each value shows the mean across runs with standard deviation in parentheses. Motion and Signal-to-Noise data are post exclusions. Pres. error = experimenter error in trial presentation. EP = Episodic Projection; ToM = Theory of Mind; FIX = Fixation.

**Table 3.**
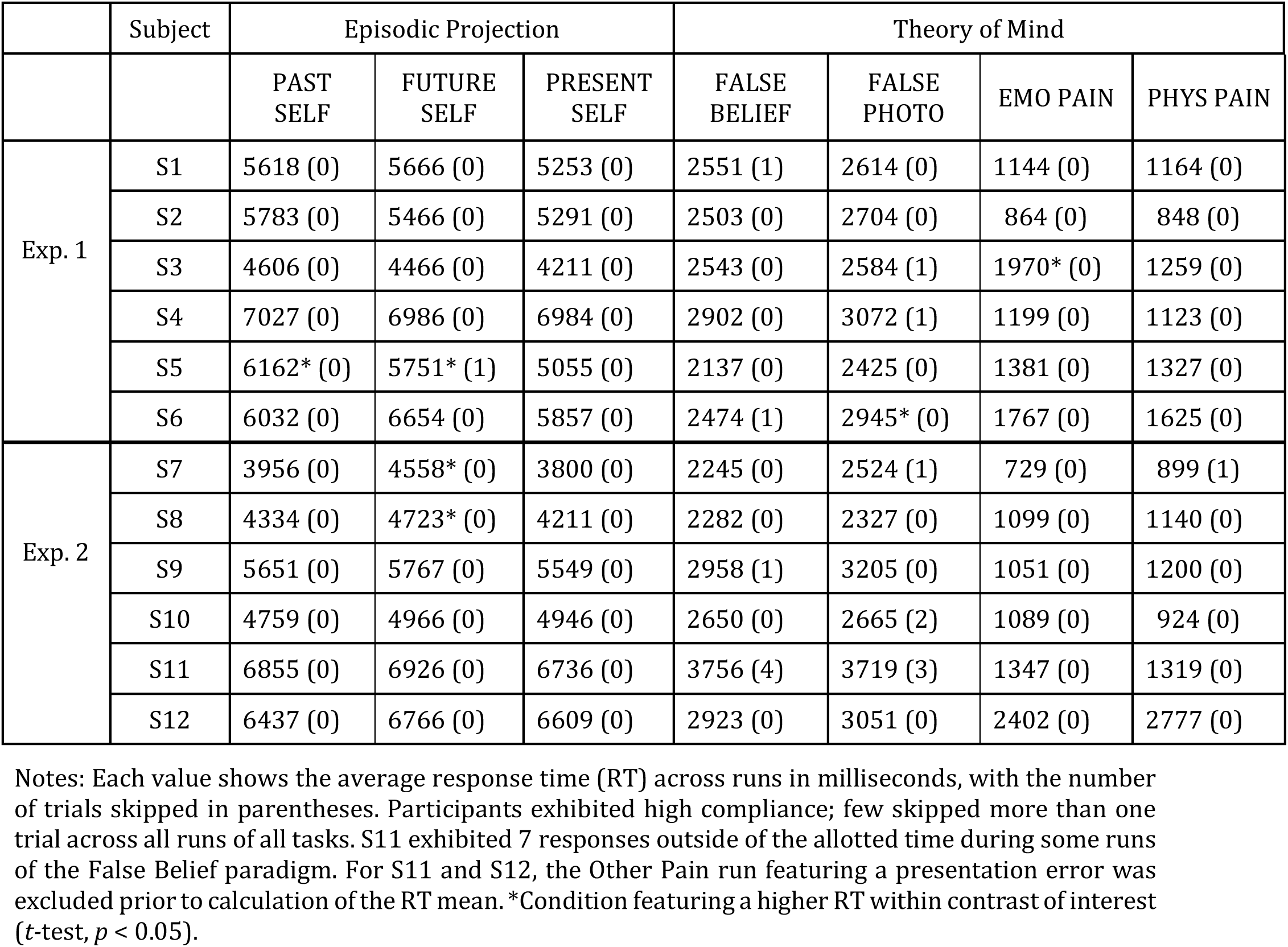
Behavioral performance across tasks.

### Within-Subject Data Processing and Template Alignment

A custom analysis pipeline for individualized data processing (“iProc”) expanded upon previous methods (e.g., Braga and Buckner 2017; Poldrack et al. 2015; Yeo et al. 2011; described in detail in Braga et al. 2019). To minimize spatial blurring, data were registered to an isotropic 1mm native space anatomical template through a single interpolation that combined four registration matrices. Prior to the calculation of these matrices, for each BOLD run, the first 12 volumes were discarded for T1 equilibration. (1) Within each run, each volume was motion-corrected to the middle volume using linear registration and 12 degrees of freedom (dof; using MCFLIRT, FSL v5.0.4; Jenkinson et al. 2002; Smith et al. 2004). (2) Each run’s middle volume was then field-map unwarped using a field map acquired during the same scanning session (using FUGUE, FSL v4.0.3; Jenkinson et al. 2004). (3) Field-map unwarped middle volumes from each run were then registered to a subject-specific, mean-BOLD template image. The mean-BOLD template was constructed iteratively. First, a single volume was selected as a temporary target: the field-map unwarped middle volume from the run acquired closest to the field map during the first scanning session. This target image was upsampled to 1.2mm isotropic resolution to optimize alignment precision, and all runs’ field-map unwarped middle volumes were registered to this upsampled target. A mean was then taken to reduce bias toward a single run. The mean image acted as a subject-specific, field-map unwarped and upsampled mean-BOLD template image, to which the field-map unwarped middle volumes from each run were linearly registered (12 dof; using FLIRT, FSL v5.0.4; Jenkinsonr & Smith 2001). (4) The mean-BOLD template was then registered to a native space template (6 DOF; using boundary-based registration; Greve and Fischl 2009). The native space template was one of the subject’s T1 structural images, upsampled to 1mm isotropic resolution and deemed as having a robust estimate of the pial and white-matter boundaries (as constructed by FreeSurfer recon-all; Fischl et al. 1999).

In this fashion, matrix calculations progressed from run- and session-specific corrections, to cross-session registration, to alignment to a template space. During matrix calculations, data were checked for registration errors. Matrices 1-4 were combined to allow for a single interpolation from each raw volume of BOLD data to the individual’s native space template. The iProc pipeline thus allowed for high-resolution and robustly aligned BOLD data, with minimal interpolation and signal loss, output to both the 1mm native space (through a single interpolation) and to the fsaverage6 cortical surface (to which native space data were resampled).

### Within-Subject Functional Connectivity Network Analysis

Functional connectivity analysis targeted identification of Networks A and B within each individual (as in Braga and Buckner 2017; Braga et al. 2019). Nuisance variables, representing a combination of 6 motion parameters, as well as whole-brain, ventricular and deep cerebral white matter mean signals and their temporal derivatives, were regressed. The residual BOLD data were then bandpass filtered at 0.01 to 0.1Hz. Preprocessed data were sampled to the fsaverage6 surface mesh (featuring 40,962 vertices per hemisphere; Fischl et al. 1999) using trilinear interpolation, then smoothed along the surface using a 2mm full-width-at-half-maximum (FWHM) Gaussian kernel. A bespoke cortical surface template (Braga and Buckner 2017) was employed to optimize interactive visualization of the surface data at this vertex density using Connectome Workbench software (wb_view; Glasser et al. 2013; Marcus et al. 2011). After projection to the surface, networks were identified within each subject using both seed-based and k-means parcellation techniques.

For seed-based network identification, mirroring the procedures outlined by Braga and Buckner (2017), a cross-correlation matrix was created for each fixation run by computing the Pearson’s *r* correlation values between time series at each vertex, and the matrices across runs were averaged together (after normalization) for a single subject. A correlation threshold of 0.2 was used, and primary seed vertices were hand-selected from the lateral PFC. Functional connectivity maps, featuring distributed sets of cortical regions with correlated patterns of signal fluctuations, were viewed for each selected seed, and optimal seeds were chosen for isolating Networks A and B (see also Braga et al. 2019).

For k-means parcellation, 17 clusters were specified (e.g., Yeo et al. 2011) to estimate whole-brain, within-subject network organization. For k-means clustering, time series data were input to the k-means algorithm following z-normalization and concatenation across runs, within a subject. Clustering was done using the MATLAB v2015b *kmeans* function, with default parameters (1 random initialization, 100 iterations, squared Euclidean distance metric). Vertices along the medial wall were removed prior to calculating the parcellation (e.g., Fig. 1A). Networks A and B were identified in the whole-brain k-means output based on referential features of each network’s anatomical distribution (described in detail in Braga and Buckner 2017; Braga et al. 2019).

As will be shown, seed-based and k-means parcellation yielded similar (but not identical) network estimates. In order to keep procedures consistent and unbiased for our primary goal of functional dissociation, we utilized the automated k-means definition of networks. However, it is important to present and contrast network definitions across both strategies as a reminder that detailed features of topography are sensitive to the exact methods employed.

### Within-Subject Task Analysis

For each run of the Episodic Projection and ToM tasks, the whole-brain signal was regressed from native-space data. Data were then resampled to the fsaverage6 cortical surface mesh (Fischl et al. 1999; Braga and Buckner 2017), smoothed using a 2mm FWHM kernel, and input to run-specific general linear models (created using FEAT; FSL version 5.0.4). All conditions were included in each model design, even those not relevant to the contrasts of interest. A canonical double-gamma hemodynamic response function (HRF) convolution was applied. Time derivatives were also included to account for HRF variability across brain regions, and data were high-pass filtered with a 100s (0.01Hz) cut-off to remove low-frequency drifts within each run. *z*-weighted outputs for each run of each task were averaged to create a single cross-session map for each contrast of interest. The mean *z*-maps provided a best estimate of the targeted Episodic Projection and ToM contrasts within each individual.

### Within-Subject Functional Double-Dissociation

The critical test of functional dissociation was to determine if the task domains exhibited differential BOLD response within the spatially separate Networks A and B. This test was achieved by quantifying to what degree the voxel response distributions within each network differed as a function of task domain (Fig. 1). A shift in the response distribution within a network would provide positive evidence for dissociation within an individual. A double dissociation would arise if the response distributions shifted in opposite directions between the two networks. Generalization of the double dissociation would be supported by replicating the opposing effects across multiple individuals. Thus, while within-individual analyses differ in many ways from more common random-effects strategies of group analyses, specific hypothesis-directed analyses were adopted to formalize the predicted double-dissociation as well as to establish whether any patterns were limited to specific individuals or were more general, shared features (see Shallice 1988 for theoretical motivation).

Specifically, after identifying Networks A and B within an individual using k-means parcellation (Fig. 1A), borders of each network were defined (Fig. 1B) and overlaid upon each domain-specific mean contrast map (Fig. 1C). Using Connectome Workbench tools (wb_command; Marcus et al. 2011), *z*-values from all vertices within the bounds of either Network A or B were extracted from each task domain’s unthresholded contrast map. For each network, distributions of *z*-values associated with each task domain were then plotted and compared. Cohen’s *d* effect sizes were calculated as a descriptive statistic to quantify differences between distributions.

This method was first employed to analyze network recruitment by task domain across Networks A and B in their entirety, and was then extended to examine recruitment more selectively for Network A regions in ventral PCC / RSC and PHC. For this more focused analysis, regions of interest approximating the whole of the ventral PCC / RSC and posterior PHC were defined on the fsaverage6 surface template. For each individual and each domain-specific mean contrast map, z-values were then extracted from vertices that fell within the bounds of both the region of interest and the individual’s Network A. In this way, the restriction to subzones was applied uniformly across subjects, but analyses were constrained to individual-specific network regions (see Figs. 4 and 11; e.g., Fedorenko et al. 2010). Cohen’s *d* effect sizes were then calculated to quantify distribution differences.

**Figure 5.**
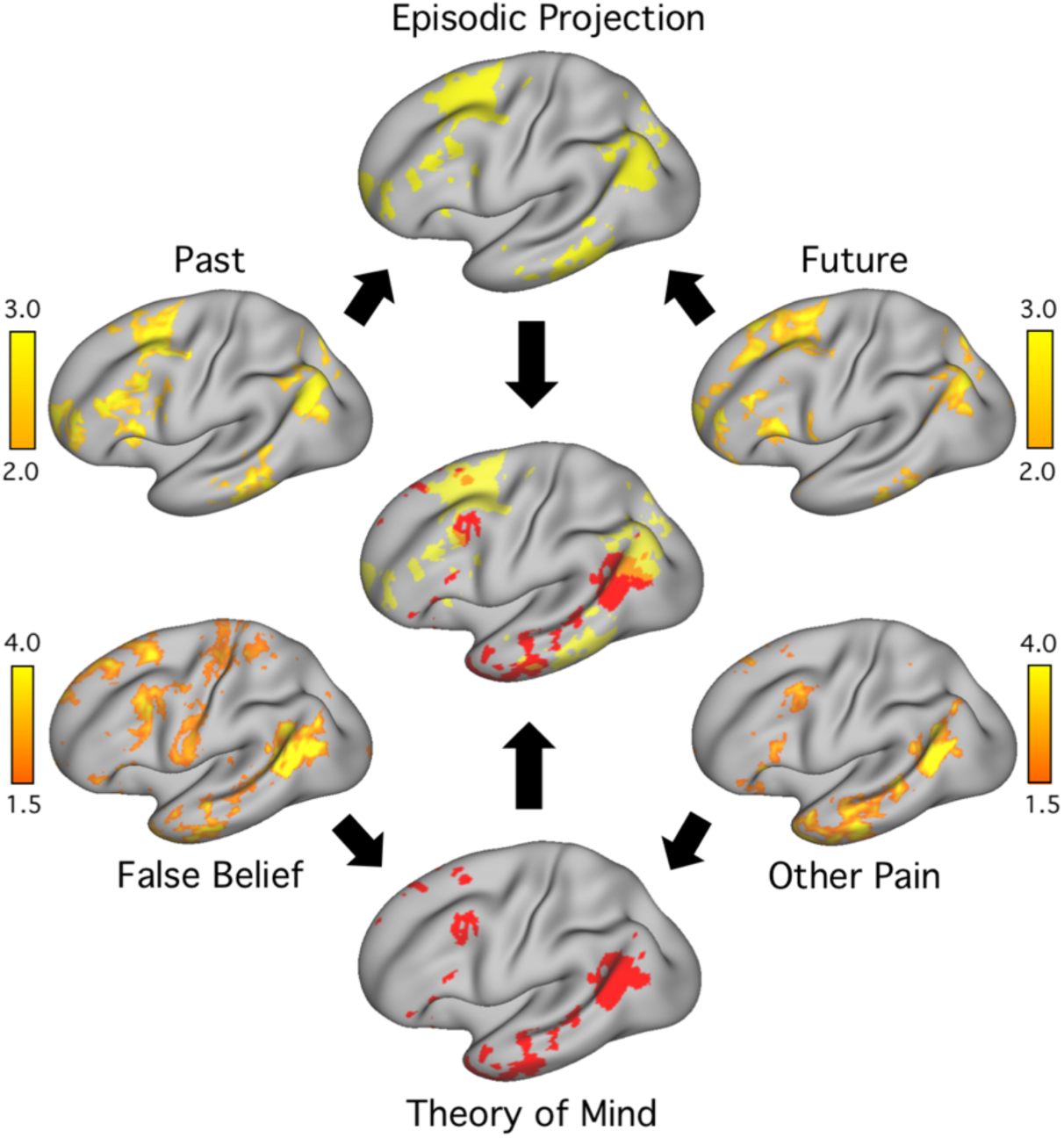
Procedure for visualizing task response patterns within individuals. Within each individual, the separate task contrasts were first estimated. Past: PAST SELF versus PRESENT SELF; Future: FUTURE SELF versus PRESENT SELF, within the Episodic Projection domain. False Belief: FALSE BELIEF versus FALSE PHOTO; Other Pain: EMO PAIN versus PHYS PAIN within the Theory of Mind (ToM) domain. Color bars indicate z-value. Within each domain, the contrasts were averaged to yield a single best estimate. The two domain maps were then thresholded and plotted on the same brain (center image) to reveal overlap (ToM = red, Episodic Projection = yellow, Overlap = orange).

**Figures 6.**
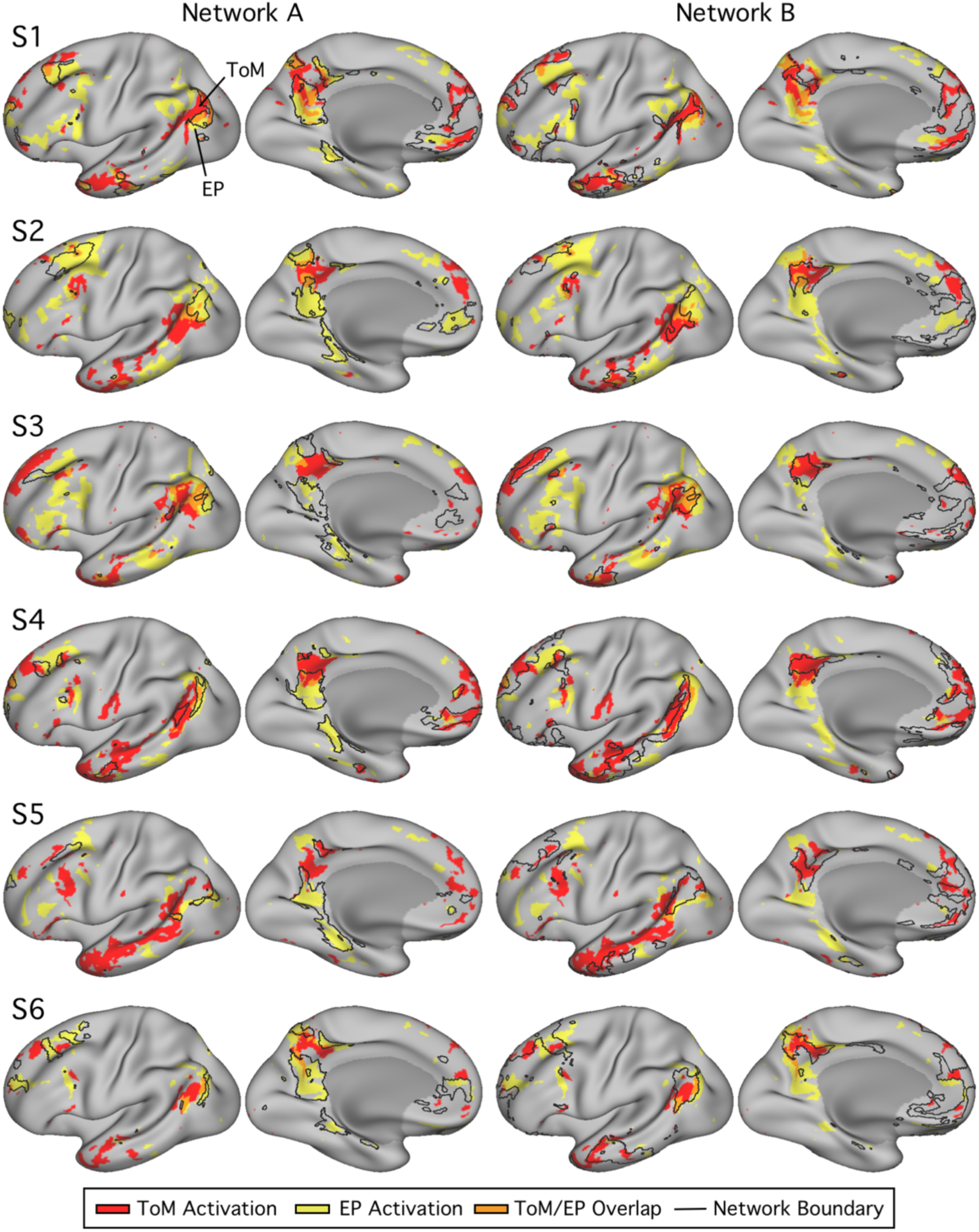
Networks A and B exhibit differential recruitment by Episodic Projection and ToM tasks across multiple cortical zones within individuals. Within each column, the lateral (left) and medial (right) surfaces of the left hemisphere are shown. The colors represent the task responses (ToM = red, Episodic Projection = yellow, Overlap = orange; see Fig. 5). For each subject, the left column displays the functional response patterns in relation to the Network A boundaries. The right column shows the same response patterns in relation to the Network B boundaries. The network boundaries are illustrated by black outlines. Episodic Projection and ToM are either partially or fully dissociable across multiple cortical regions in all subjects. The most striking double dissociations are evident in S2 and S6, where small idiosyncratic features of the differential task response patterns are predicted by the network boundaries in all zones of cortex.

**Figure 7.**
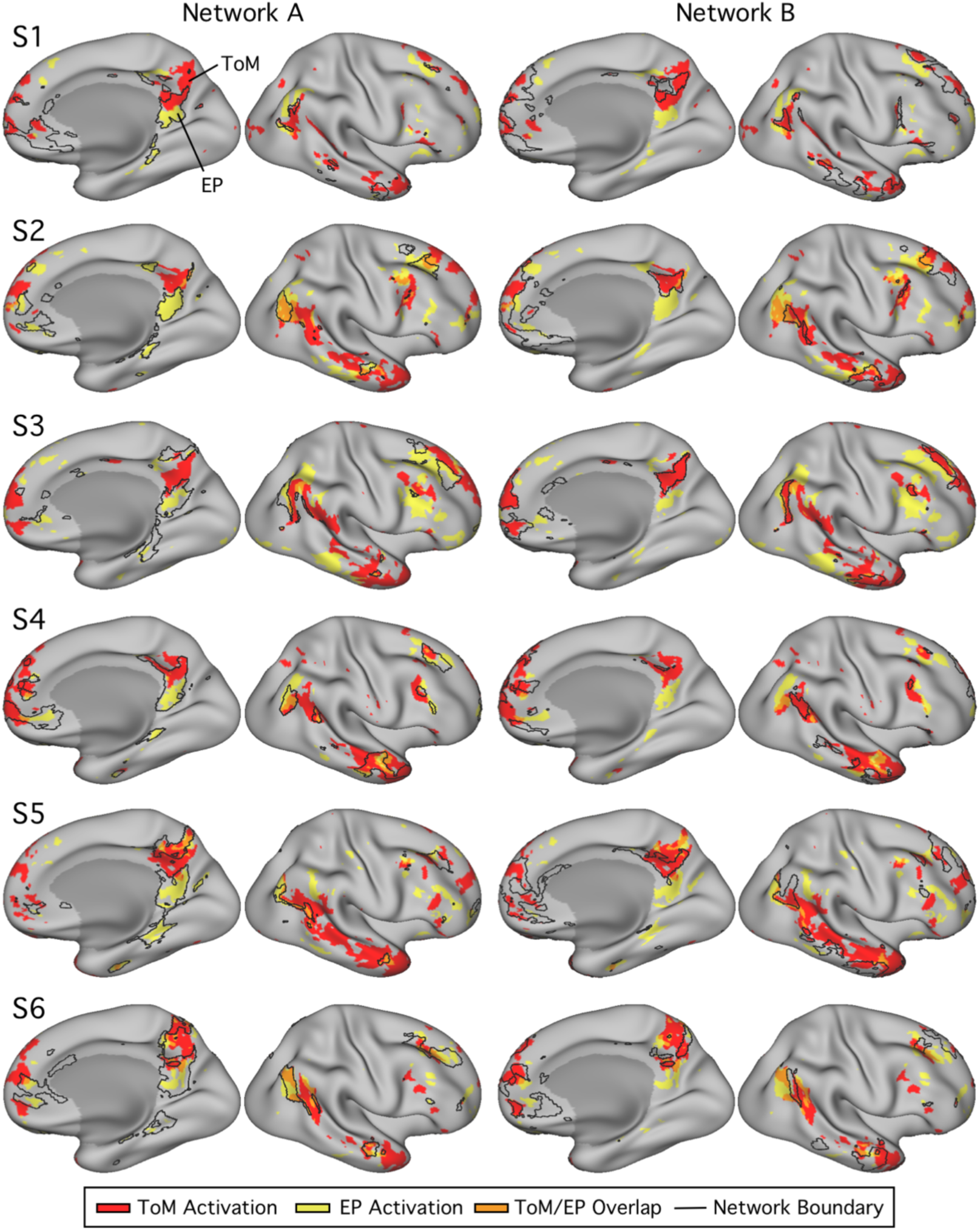
Networks A and B exhibit differential recruitment by Episodic Projection and ToM tasks in the right hemisphere. Using the same procedures as displayed in Fig. 6, the right hemisphere is displayed for each of the 6 subjects from Exp. 1. Each row shows the lateral (right) and medial (left) views of the right hemisphere for all subjects, with Network A boundaries (left columns) or Network B boundaries (right columns).

**Figure 8.**
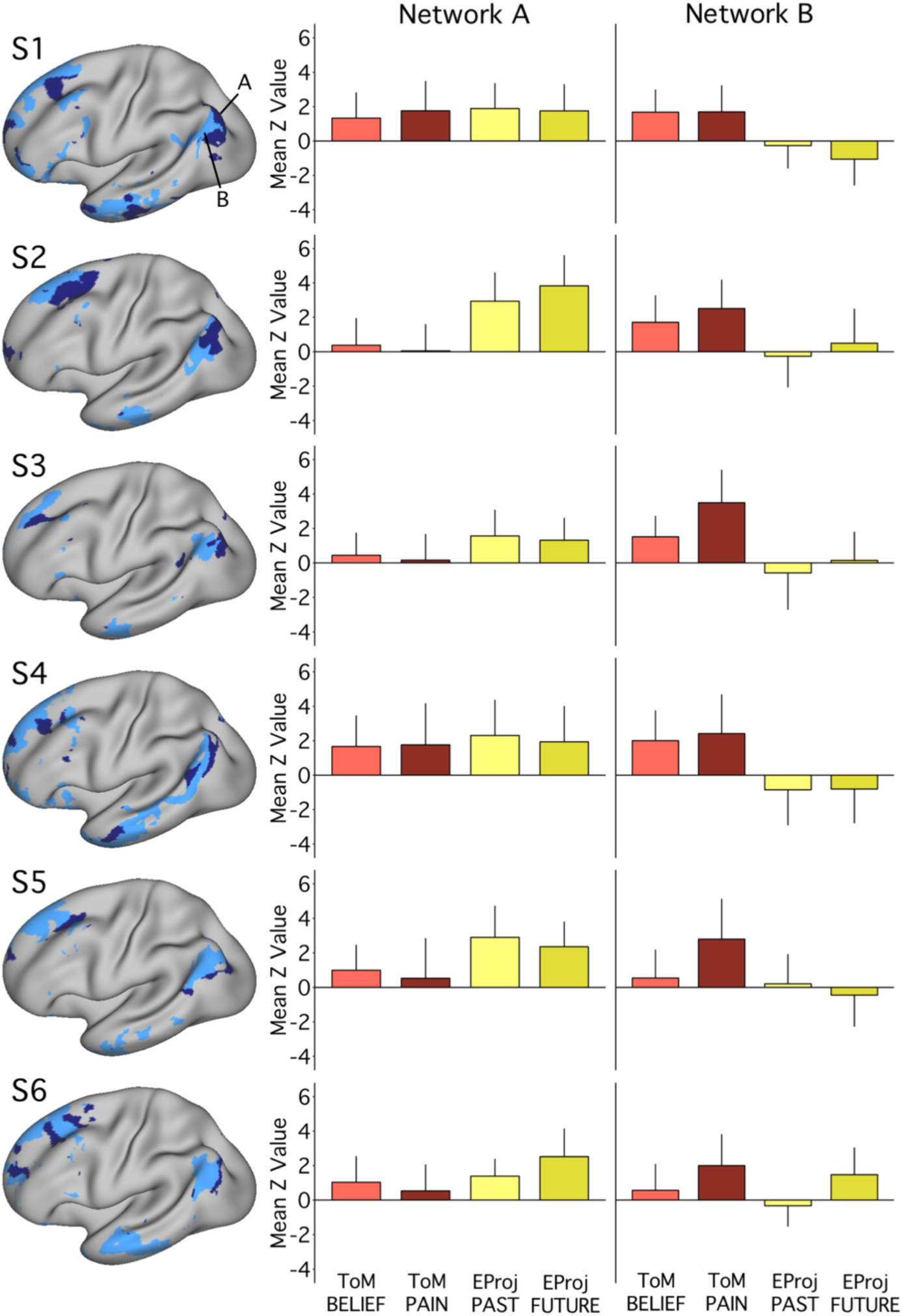
Functional dissociation of Networks A and B observed across task contrasts in Exp. 1. The left column again illustrates the spatial distributions of Network A (navy) and Network B (light blue) for each subject in Exp. 1. The bar graphs plot the functional responses (mean *z*-values and standard deviations across runs) within each network for each task-specific contrast. Tasks within a domain exhibit comparable patterns of network recruitment. Within Network B, for several individuals, the Other Pain contrast exhibits stronger recruitment than the False Belief contrast. For Network A, functional response increases for both Episodic Projection contrasts over both ToM contrasts are evident in 5 out of 6 subjects (Cohen’s *d* range = 0.08-2.25 for pairwise comparisons, with most *d* > 0.80). For Network B, increases for both ToM contrasts over both Episodic Projection contrasts are also evident in 5 out of 6 subjects (*d* range = 0.20-2.01, with most *d* > 1.20).

**Figure 9.**
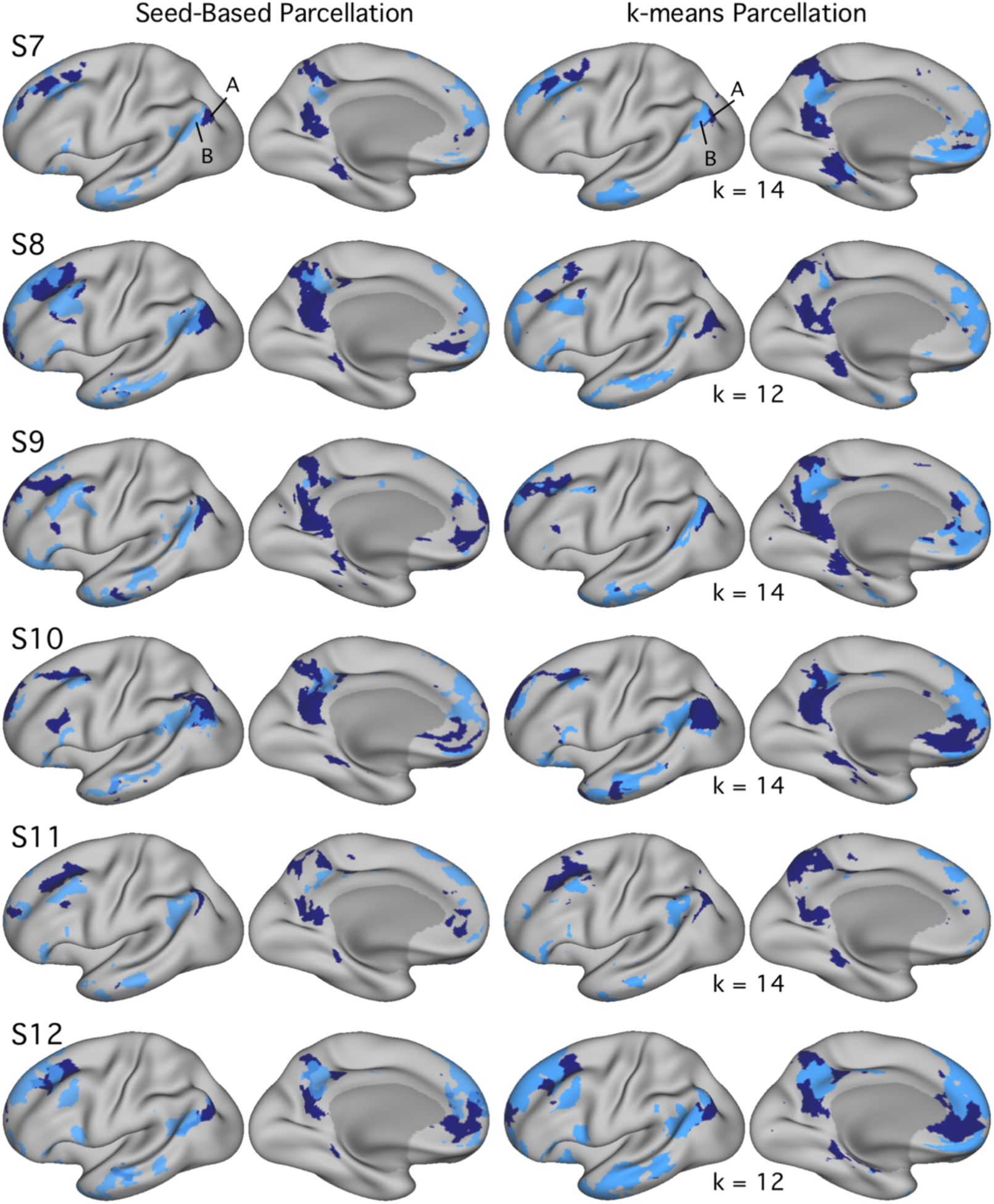
Networks A and B identified within individuals in Exp. 2 using both seed-based and k-means parcellation strategies. Estimates of Networks A and B identified using seed-based (left column) and k-means (right column) methods exhibit comparable maps within individuals in Exp. 2. In Exp. 2, *k* was set to the lowest number of clusters featuring differentiation of Networks A and B, labelled for each individual. Network A appears in navy and Network B in light blue. Similar to Exp. 1 results, estimated network boundaries differ by method, but the two networks could be identified in all individuals with both methods. Seed-based maps are thresholded at *r*=0.40.

**Figure 10.**
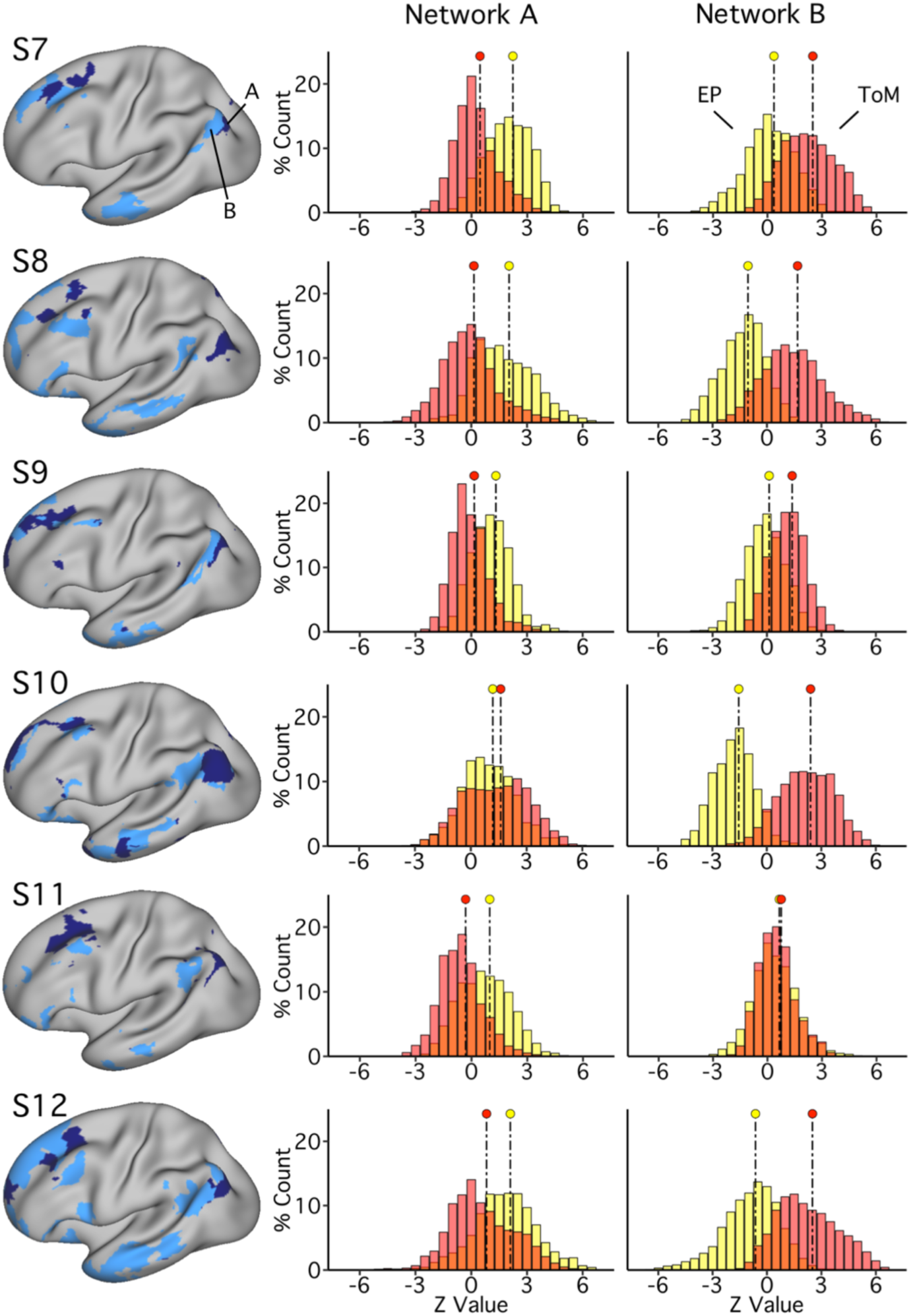
Networks A and B replicate functional dissociation within individuals in Exp. 2. The left column features the spatial distributions of Network A (navy) and Network B (light blue) for each subject as defined by k-means clustering. The distributions in the right columns plot the functional responses within each network for the two task domains (ToM = red, Episodic Projection = yellow, Overlap = orange). For Network A, 5 out of 6 subjects demonstrate a functional response increase for Episodic Projection over ToM contrasts (Cohen’s *d* range = 0.75-1.52). For Network B, all subjects demonstrate the opposite pattern -- an increase for ToM over Episodic Projection (*d* range = 0.09-3.01; *d* > 1.20 for all except S11). These findings replicate and support a functional double dissociation.

**Figure 11.**
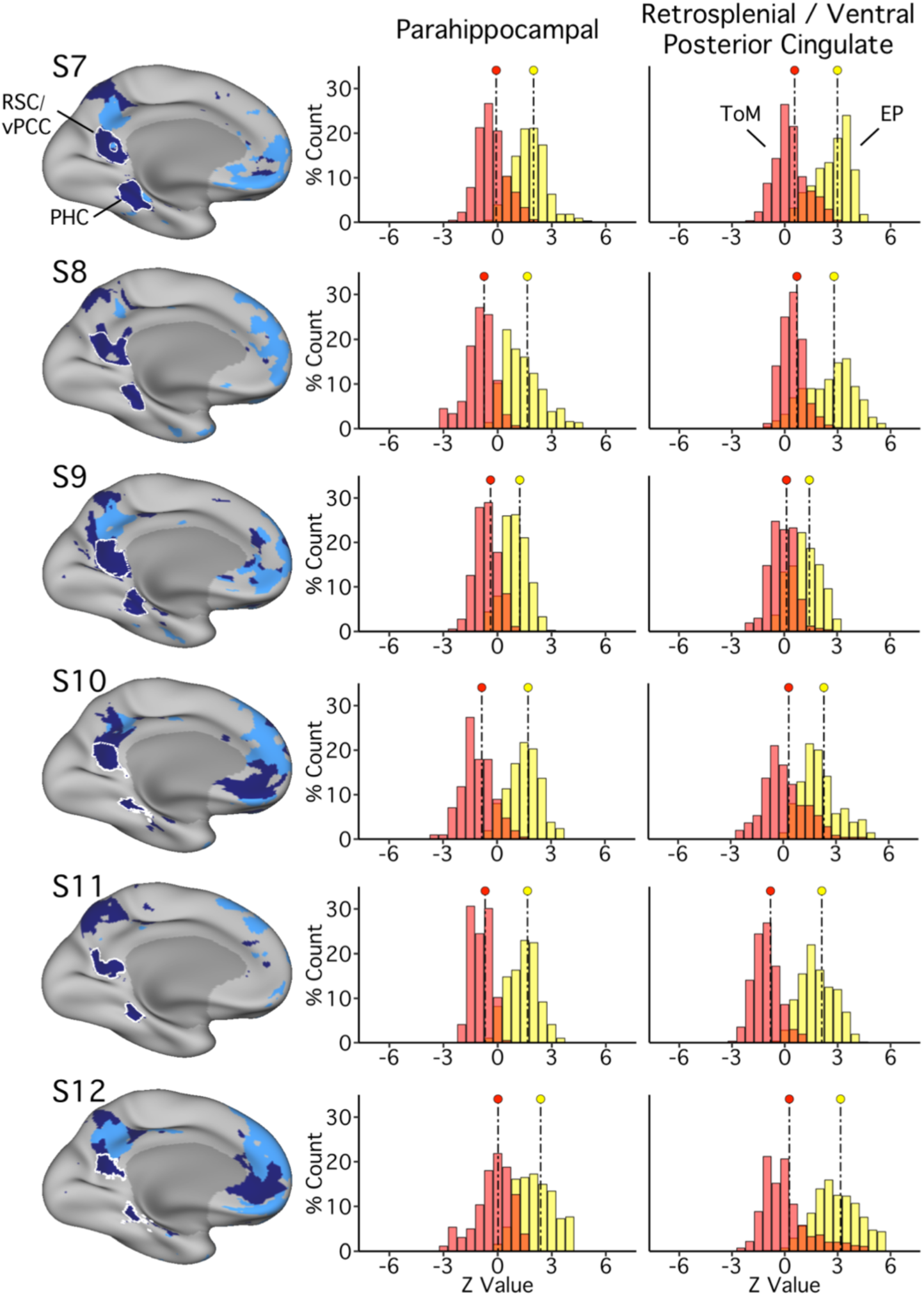
PHC and ventral PCC/RSC regions of Network A replicate functional dissociation in Exp. 2. The left column illustrates PHC and RSC / vPCC regions as defined in each subject. These regions are outlined in white and contained within Network A, as in Fig. 4. The distributions plot the functional responses within each region for the two task domains (ToM = red, Episodic Projection = yellow, Overlap = orange). For each region, in all subjects, a significant functional response increase for Episodic Projection over ToM is evident (Cohen’s *d* ranges: RSC/vPCC: 1.60-3.42; PHC: 2.35-3.39).

We chose a quantitative descriptive approach because each network features a complex correlation structure spanning spatially discontinuous regions. Methods for identifying false positives that assume spatial dependencies only between neighboring voxels fail to account for the complex nature of correlations in the present data. Here, across analyses, we present descriptive statistics and base interpretations on evidence of predicted double-dissociation patterns consistently generalizing across individuals. To alleviate any ambiguity about the robustness of the results, the entire approach and findings were replicated in Exp. 2. As will be shown, the key patterns, including the double dissociation, were evident in the vast majority of subjects and compelling in many individual subjects.

### Within-Subject Visualization of Task Response Maps

For each participant, contrast maps for each domain were visualized in relation to the independently derived functional connectivity network boundaries. Borders of Networks A and B were overlaid on an image that displayed both task domain contrast maps on the same cortical surface. The contrast maps were thresholded for visualization (with minimum *z-*values chosen between 1.5 and 2.5 for each subject). Patterns of overlap between each network and the domain-specific task responses were visualized across multiple cortical zones to clarify whether observed dissociations were carried by specific regions or reflected functional specialization across the broad distributed networks.

### Contrast-Specific Analyses

For the primary hypothesis-testing analyses described above, we collapsed across tasks within Episodic Projection and ToM domains in order to optimize our power for detecting domain-related differences between networks. Tasks were chosen *a priori* based on their likelihood to isolate component processes within each domain. A natural question to ask is: do tasks within each domain exhibit similar patterns of network recruitment? To address this question and provide a full description of the data, we conducted a *post-hoc* analysis to quantify potential distinctions between specific task exemplars, by plotting and comparing contrast-specific means. *z-*values from vertices within the bounds of each network were extracted from each separate task’s mean, unthresholded contrast map and plotted. Effect sizes were calculated to describe potential differences.

## Experiment 2 Methods: Prospective Replication

### Participants

Exp. 2 was conducted as a prospective replication of Exp. 1. An independent sample of 6 adults was recruited (*Mean* = 22.3 years (SD = 4.0), 2 males, 5 right-handed) to complete 4 scanning sessions each. All paid participants were native English speakers, were screened to exclude neurological or psychiatric illness, and provided written informed consent using procedures approved by the Institutional Review Board of Harvard University.

MRI data acquisition and processing methods, as well as most task and analysis procedures in Exp. 2, were the same as for Exp. 1; differences are detailed below.

### Fixation Runs for Intrinsic Functional Connectivity Analysis

Eleven 7m 2s BOLD fixation runs were acquired per individual (approximately 77m of data; 3 runs during each of the first three sessions, and 2 runs during the fourth).

### Task Paradigms

Exp. 2 included the same critical Episodic Projection and ToM task as in Exp. 1. Additional reference tasks were added to expand the possible contrasts involving Episodic Projection.

#### Expanded Episodic Projection Paradigm

The Episodic Projection contrasts (involving PAST SELF and FUTURE SELF) each subtracted a self-referential control (PRESENT SELF), thereby removing self-reference from the contrasts of interest. Although this was critical to the primary goal of targeting Episodic Projection as distinct from ToM, particularly given that shared processes may allow for reflecting on one’s own and others’ mental states (e.g., Lieberman 2007), contrasts featuring self-reflection *and* episodic projection might be expected to call upon both Networks A and B. To begin to address this possibility, in Exp. 2, the Episodic Projection paradigm was expanded. In addition to the three primary conditions from Exp. 1 (PAST SELF, FUTURE SELF and PRESENT SELF), the Episodic Projection paradigm in Exp. 2 included three nonpersonal conditions featuring semantic questions about the past (PAST NON-SELF), present (PRESENT NON-SELF) or future (FUTURE NON-SELF). Structured to resemble the setup of Andrews-Hanna et al. (2010), this paradigm featured a 2×3 design, with questions varying along dimensions of self-relevance (SELF versus NON-SELF) and temporal orientation (PAST, PRESENT, OR FUTURE). Each self-relevant condition, therefore, was time-matched to a nonpersonal semantic control, producing three contrasts that were not possible in Exp. 1: PAST SELF versus PAST NON-SELF, PRESENT SELF versus PRESENT NON-SELF and FUTURE SELF versus FUTURE NON-SELF.

Given prior findings that episodic future projection preferentially activates components of Network A (Andrews-Hanna et al. 2010), and that self-reflection recruits regions also associated with ToM (Andrews-Hanna et al. 2010; see also Lieberman 2007), we hypothesized that the episodic FUTURE and PAST contrasts might preferentially recruit Network A, while all three self-referential contrasts might recruit Network B.

All task conditions were intermixed in each run, to create randomized trial orders. Each 612s run included 30 trials (5 per condition), with timing matched to Exp. 1. Sentence structure, words and character length were again matched across conditions. Two runs of 30 trials were collected during each of three MRI sessions, yielding 6 total runs (180 individual trials, 30 of each type; 60m of total Episodic Projection data).

#### Theory of Mind Tasks

Conditions and stimuli were the same as those in Exp. 1 for both ToM tasks. Four 5m runs of each task were acquired per individual across three MRI sessions (40m of total ToM data).

### Exclusion Criteria and Quality Control

Exclusion criteria were carried forward from Exp. 1 (see Table 2), but a slightly more stringent maximum motion cutoff of 1.8mm was used. Across subjects, 4 of 150 runs were excluded in Exp. 2 for motion. For each subject, the mean maximum motion displacement in included data was less than 1mm. Behavioral performance across tasks was also examined for each participant, with no runs excluded based on performance metrics (see Exp. 2 Results and Table 3 for details). One individual in Exp. 2 (S9) discontinued participation after two sessions, resulting in a total of 6 fixation runs (approximately 42m), 6 ToM task runs (30m) and 4 Episodic Projection task runs (40m) for this subject.

### Within-Subject Data Processing and Template Alignment

iProc was used for data processing and template alignment for each individual in Exp. 2, with procedures that matched those from Exp. 1. Of note, data from two participants (S7 and S8) were processed following an upgrade to both the Freesurfer version (from a beta release of v6 to published v6.0.0) and system software (from centOS6 to centOS7). The iProc procedures, commands and software versions were otherwise identical across experiments and subjects. Extensive testing revealed minimal differences between data pre-processed using iProc on the original versus upgraded systems, and since all analyses were conducted within-subjects, any small residual differences would not influence results.

### Prospective Replication of the Functional Double Dissociation

Functional connectivity procedures were preserved from Exp. 1. k-means estimates were calculated for solutions between *k*=10 and *k=*17; the solution with the fewest clusters that also featured differentiation between Networks A and B was chosen for each individual prior to examination of any task data.

In two individuals (S8 and S11; Fig. 9), although the clustering analysis revealed PHC-linked Network A and TPJ-linked Network B, the regions of these networks were not always side-by-side as was found in other subjects. Instead, a third cluster was observed, occupying regions at the spatial intersection of Networks A and B. It is possible that this third cluster was a result of sufficient blurring between Networks A and B at multiple cortical sites. Given that our hypothesis regarded PHC- and TPJ-linked distributed networks, the third cluster was not utilized for further analysis.

All procedures used to obtain the results for Exp. 1, to obtain and quantify task contrast response distributions, were then applied, without any iterative adjustments, to the data from Exp. 2, even including how the data distributions and maps were plotted.

### Exploratory Analysis of Network Recruitment for Self-Referential Processing

Beyond replication of Exp. 1, the Episodic Projection paradigm in Exp. 2 also allowed for exploratory analysis of network recruitment across three new self-referential contrasts, each targeting a separate timeframe (PAST, PRESENT, or FUTURE). Within-subject task analysis procedures (e.g., whole-brain signal regression, surface resampling, and run-specific general linear models) mirrored those described for the primary Episodic Projection and ToM task contrasts. Similar to the whole-brain and region-specific distributional analyses described, *z-*values from vertices within the bounds of each network were extracted from each of the three unthresholded mean contrast maps. The *z-*values were then plotted and compared, and effect sizes quantified differences between contrast distributions. A positive shift in response distributions across contrasts requiring episodic judgments (those featuring PAST and FUTURE conditions) was predicted for Network A; a positive shift, across all three self-referential contrasts, was predicted for Network B.

## Experiment 1 Results

### Behavioral Performance

Participants missed a maximum of a single trial each, across all runs of all task contrasts (see Table 3). RTs were largely similar across the ToM contrasts of interest in each task (FALSE BELIEF versus FALSE PHOTO, *p* > 0.05 for all within-subject *t*-tests except S6; EMO PAIN versus PHYS PAIN, all *p* > 0.05 except S3), as well as across the Episodic Projection contrasts (FUTURE SELF versus PRESENT SELF, *p* > 0.05 except S5; PAST SELF versus PRESENT SELF, *p* > 0.05 except S5). For each individual, all acquired task data were included in subsequent analyses.

### Parallel Interdigitated Networks Are Identified Within Individuals

Networks A and B were identified within individual subjects using seed-based and k-means parcellation strategies (Fig. 2). Replicating Braga and Buckner (2017; see also Braga et al. 2019), distinct networks were found for all subjects, using manually-placed seed regions in left prefrontal cortex. Given that seed-based methods introduce bias near the seed regions and are sensitive to manual placement, a data-driven approach to network identification was also employed: k-means parcellation. Within-subject estimates of Networks A and B could be identified that were highly similar to the seed-based networks, thus establishing that the broad distinction between the networks is not reliant on a specific analysis strategy (Fig. 2; see also Braga et al. 2019).

In all individuals, the two parallel, interwoven networks spanned frontal, temporal and parietal cortices and exhibited ‘diagnostic’ features (Braga et al. 2019): Network A showed connectivity to posterior PHC, largely absent in B, while Network B included a more rostral portion of the IPL extending into the TPJ. In addition, Network A included representation at or near RSC, ventral to a region of Network B along the posterior midline. Network B featured a small region of the medial PFC ventral to that of Network A, occupied a greater portion of lateral temporal cortex than Network A, and, in most cases, included a region along the lateral inferior frontal cortex, which was absent or smaller and more dorsally-positioned in Network A.

While the majority of network details generalized across individuals with both network estimation methods, there were some details that differed among the subjects and between the methods. In several participants, for example, whether defined through seed-based or k-means parcellation, Network A showed minimal representation in left lateral temporal cortex, a region included within the original description of Network A (Braga and Buckner 2017). The Network A representation in lateral temporal cortex was robust in several of the individuals, suggesting that this is either a region of particularly complex geometry or additional data may be necessary for full specification of lateral temporal features in all individuals. Idiosyncratic patterns of signal dropout may also obscure this region in some individuals.

In relation to network selection method, while the general patterns were similar using either approach, the exact boundaries differed between the methods. This is expected to some degree because k-means parcellation forces a winner-takes-all separation. That the outer boundary and specific details of a region’s extent and location differed between methods is a reminder that each method has different sensitivities and should be considered an approximation of the true underlying organization. These details should not detract from the broader observation that the two networks could be identified in all individuals, with both methods (Fig. 2).

Given our goal of formally testing for functional dissociation between the networks, we carried forward the unbiased network estimates from the k-means parcellation.

### Networks A and B Show Functional Dissociation Within Individuals

Our primary hypothesis was that Episodic Projection would preferentially recruit PHC-associated Network A and ToM would preferentially recruit TPJ-associated Network B. To quantify each network’s functional recruitment, within a subject, the distributions of BOLD responses across vertices for each task domain were directly contrasted for each network (Fig. 3).

The dissociation was robust; all 6 subjects exhibited the predicted pattern of differential network recruitment across 12 out of 12 distribution comparisons. Differences between distributions were strong across individuals for Network B (6 out of 6 subjects; Cohen’s *d* range: 0.51-1.76) and present also for Network A, although weaker in 2 subjects (Cohen’s *d* range: 0.88-2.03 for 4 subjects; *d* = 0.19 for S1 and *d* = 0.20 for S4). Initial quantitative analysis of functional recruitment by task domains thus revealed evidence for a double dissociation in all 6 subjects. Since 2 subjects exhibited weaker differentiation for Network A, these results motivated further analysis of Network A, focusing on two component regions that differentiate this network most strongly (Braga and Buckner 2017).

### Parahippocampal and Retrosplenial / Ventral Posterior Cingulate Regions of Network A Show Strong Functional Dissociation

Initial findings revealed variation across subjects in Network A recruitment by task domain. *Post hoc* analyses were conducted for the two regions in Network A most closely linked to the MTL memory system, allowing a robust test for dissociation, albeit in a subset of Network A regions. Procedures identical to those used in the primary distribution analysis (Fig. 1) were limited to vertices within a ventral PCC / RSC region and separately within a PHC region (Fig. 4). All 6 subjects exhibited robust preferential recruitment of ventral PCC / RSC and PHC regions of Network A by Episodic Projection tasks over ToM tasks (Cohen’s *d* ranges – ventral PCC / RSC: 0.54-4.43; PHC: 1.67-3.24). These results further support dissociable functional responses of Networks A and B but leave open the possibility that the double dissociation does not pertain to all of the distributed zones of cortex.

### Preferential Recruitment is Evident Across Multiple Cortical Zones

The analyses above provided evidence for differential network recruitment by tasks from Episodic Projection and ToM domains. However, the described quantitative results could align either with fully distinct networks or with previously proposed network configurations featuring dissociable network features *and* shared “core” regions (Andrews-Hanna et al. 2010). That is, the described dissociation results could be carried out by *subsets* of network regions. To explore whether recruitment differences were limited to specific cortical zones or were distributed, the differential task domain responses were visualized for each individual (Fig. 5) in relation to the estimates of Networks A and B.

Figure 6 displays the full task contrast maps for each individual in Exp. 1, in relation to Network A and B boundaries. The first broad observation is that the Episodic Projection and ToM contrast maps have non-overlapping components in most zones of cortex. It is not simply that certain zones show spatial functional separation, such as TPJ versus more caudal IPL, but rather that spatial separation could be observed in medial prefrontal, lateral temporal, and dorsolateral prefrontal cortices, as well as in posteromedial cortex. Thus, at a map level within individuals, the two task domains display broad functional separation across the cortex.

The second critical observation is that the functional connectivity estimates of Networks A and B within each individual predict many idiosyncratic features of the differential patterns of task response. This was not always the case in all subjects in all zones (e.g., lateral temporal cortex and medial PFC displayed some of the most complexity), but it was the case often and clearly in some individuals (e.g., S2 and S6).

Of particular note, task responses in Networks A and B showed clear spatial separation in the zones previously considered “core” regions along the cortical midline (Andrews-Hanna et al. 2010). For example, in several subjects the posteromedial representation of Network A included a triad of distinct regions: a region near ventral PCC/ RSC, as well as distinct dorsocaudal and dorsorostral regions. The Network B region of response was most typically in between these three Network A regions. In multiple cases, the differential task response followed this idiosyncratic anatomy, which is a reproducible hallmark feature of network interdigitation along the posterior midline, when visualized in surface representation or in the volume (Braga et al. 2019).

Overall, evidence from both quantitative analysis and topographic visualization suggests a functional dissociation of Networks A and B, with Network A preferentially recruited for the task contrasts isolating processes related to Episodic Projection and Network B for task contrasts isolating processes related to ToM. The dissociation does not appear to be limited to one zone of cortex but rather, as is evident in the clearest cases, reflects functional specialization across the full extent of the distributed networks, across both hemispheres (Fig. 7).

### Functional Dissociation of Networks A and B Observed Across Task Contrasts

For all analyses above we collapsed across tasks within the Episodic Projection and ToM domains, toward the primary goal of detecting domain-related distinctions in network recruitment. This was the *a priori* design of the study. As a final analysis, we examined differences across each specific task contrast.

Contrast-specific recruitment of Networks A and B (Fig. 8) bolstered the findings from the pooled results (see Fig. 3). Within Network A, for example, all subjects showed similar recruitment across ToM tasks (Cohen’s *d* range = 0.05-0.33), and 4 subjects showed similar mean recruitment across Episodic Projection tasks (*d*=0.09-0.33). For S2 and S6, the Episodic Projection tasks exhibited greater differences (*d*=0.52 for S2; *d=*0.83 for S6, FUTURE > PAST), but for these individuals and three others (S3, S4 and S5), both Episodic Projection contrasts still exhibited higher mean *z-*values than both ToM contrasts within Network A (*d* range = 0.08-2.25 for pairwise comparisons; most *d* > 0.80). Individuals featuring more ambiguous Network A distributions (see S1 and S4 in Fig. 3), exhibited similar recruitment for both ToM tasks, suggesting a subject difference rather than a task difference. Across participants, no single task contrast appeared to drive the Network A results.

Within Network B, there was evidence for a systematic task difference in the ToM domain. All 6 subjects showed a greater response for the ToM OTHER PAIN contrast relative to the FALSE BELIEF contrast. Still, even with this difference, the ToM contrasts, barring one exception, showed higher responses within Network B than either of the Episodic Projection task contrasts (Cohen’s *d* range = 0.20-2.01; most *d* > 1.20). Only S6 had a ToM mean *z-*value (for FALSE BELIEF) less than the mean of a single Episodic Projection task contrast (FUTURE). Thus, either of the two ToM task contrasts would still allow the critical double-dissociation between Networks A and B to be detected.

## Experiment 2 Results

Data collection and analyses of Exp. 2 were conducted prospectively after the results of Exp 1. were known and all analysis procedures, down to the details of exactly how results would be plotted, were settled. Thus, the findings should be considered a prospective replication.

### Behavioral Performance

Table 3 presents behavioral performance. Most participants responded to all trials, but S11 exhibited multiple skipped trials during the False Belief task. Behavioral monitoring revealed that S11 responded to each trial of this task, but not always within the timeframe necessary for recording; these data were therefore included in subsequent analysis. One ToM run (from the Other Pain task) was excluded from each of two participants (S11 and S12) due to an error in trial presentation that resulted in repeated trials.

RTs were again largely similar across the ToM contrasts of interest in each task (FALSE BELIEF versus FALSE PHOTO, *p* > 0.05 for all within-subject *t*-tests; EMO PAIN versus PHYS PAIN, all *p* > 0.05), as well as across the Episodic Projection contrasts (FUTURE SELF versus PRESENT SELF, *p* > 0.05 except S7, S8; PAST SELF versus PRESENT SELF, all *p* > 0.05). Behavioral performance metrics thus supported compliance across tasks by all individuals.

### Parallel Interdigitated Networks Replicate Across Additional Individuals

Networks A and B were again identified using both seed-based and k-means parcellation strategies (Fig. 9). The parallel networks were found to be interdigitated and distributed across frontal, temporal and parietal regions with the diagnostic spatial features for the networks again largely observed across individuals (Braga et al. 2019): Network A exhibited connectivity to posterior PHC (absent in most individuals’ estimates of Network B), and Network B included a portion of the IPL near to the TPJ, rostral to a region of Network A. Network A also showed a more ventral representation along the posterior midline, extending to or near RSC; Network B included a region ventral to Network A regions in medial PFC.

For some individuals, as in Exp. 1, Network A as estimated by k-means showed minimal representation in left lateral temporal cortex. For a few individuals (e.g., S8 and S11), the medial PFC showed smaller Network A representations and posteromedial cortex showed smaller Network B representations in the k-means than in the seed-based estimates. Medial regions were difficult to parse in group-averaged data (Andrews-Hanna et al. 2010), and complexity in these regions appears difficult to resolve within some individuals, as well. For half of the individuals (S7, S9, S11), the k-means estimates also lacked the typically strong representation of Network B along the lateral inferior frontal cortex. These details highlight that both k-means and seed-based methods produce only approximations of network organization. Despite this, overall, the two networks could be identified in all individuals, with both methods (Fig. 9).

For planned analyses in Exp. 2, we carried forward the unbiased k-means network estimates for each individual.

### Networks A and B Replicate Functional Dissociation Across Additional Individuals

Functional recruitment was quantified for each task domain within each network (Fig. 10). Results again strongly supported functional dissociation for both Network A (5 out 6 subjects; Cohen’s *d* range: 0.75-1.52) and Network B (all subjects; Cohen’s *d* range: 0.09-3.01; *d* > 1.20 for all except S11). The predicted double dissociation pattern was seen clearly in most subjects. Distinctions between the networks were most notable in a subset of individuals (e.g., S7, S8 and S12).

### Robust Functional Dissociation Replicates for Parahippocampal and Retrosplenial / Ventral Posterior Cingulate Regions of Network A

Region-specific explorations for the ventral PCC / RSC and PHC regions of Network A again demonstrated robust dissociation (Fig. 11). For all 6 individuals the region-specific analysis revealed preferential recruitment of both ventral PCC / RSC and PHC by Episodic Projection tasks over those requiring ToM (Cohen’s *d* ranges – ventral PCC / RSC: 1.60-3.42; PHC: 2.35-3.39).

### Preferential Task Recruitment Across Multiple Cortical Zones Replicates

Contrast maps revealed considerably non-overlapping response patterns between task domains (Fig. 12). Estimates of Networks A and B overlapped with task patterns in most subjects and clearly in a subset of this independent sample (e.g., S7 and S12), with Network A better predicting the Episodic Projection map and Network B the map for ToM.

**Figures 12.**
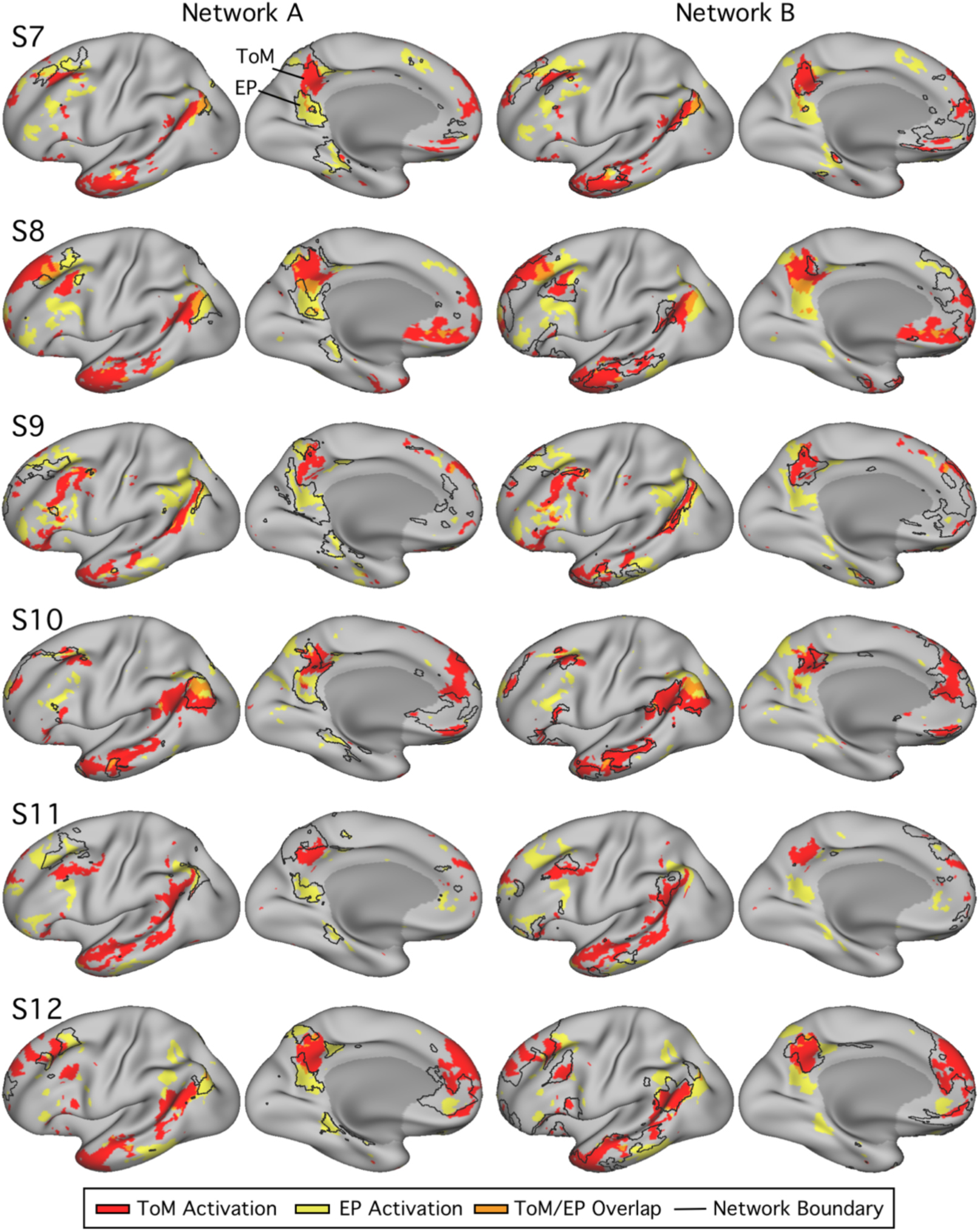
Differential recruitment of Networks A and B by Episodic Projection and ToM tasks, across multiple cortical zones, replicates within additional individuals. Within each column, the lateral (left) and medial (right) surfaces of the left hemisphere are shown. The colors represent the task responses (ToM = red, Episodic Projection = yellow, Overlap = orange; see Fig. 5). For each subject, the left column displays the functional response patterns in relation to the Network A boundaries. The right column shows the same response patterns in relation to the Network B boundaries, with boundaries shown as black outlines. Episodic Projection and ToM are either partially or fully dissociable across multiple cortical regions in all subjects in Exp. 2, replicating results from Exp. 1. The most striking double dissociations are evident in S7 and S12, where even idiosyncratic features of the differential task response patterns are predicted by the network boundaries in all cortical zones.

Of particular note, although nearly all individuals’ network estimates exhibited expected, diagnostic features (Braga et al. 2019), as an exception, the Network B estimate of S7 included a portion of posterior PHC, typically present only in Network A. As revealed in Figure 12, the task maps from each domain exhibited high similarity to network boundaries in this region. The portion of PHC attributed to Network B surrounded a region of ToM activation, whereas an adjacent portion of PHC, attributed to Network A, showed preferential response during the Episodic Projection contrast. These features would likely be lost in group-averaged data.

Task patterns along the posterior midline exhibited spatial separations that, in the clearest individuals, were well-aligned to the boundaries of Networks A and B (previously considered a “core” region of the DN; Andrews-Hanna et al. 2010). In multiple individuals, the posteromedial representation of Network A again included a triad of regions, each exhibiting preferential overlap with the Episodic Projection contrast map, while an interwoven region of Network B exhibited preferential overlap with the contrast map for ToM. Spatial alignment between networks and contrast map predictions was not limited to a specific cortical zone (e.g., see the medial and lateral PFC in S12, Fig. 12) or hemisphere (Fig. 13).

**Figure 13.**
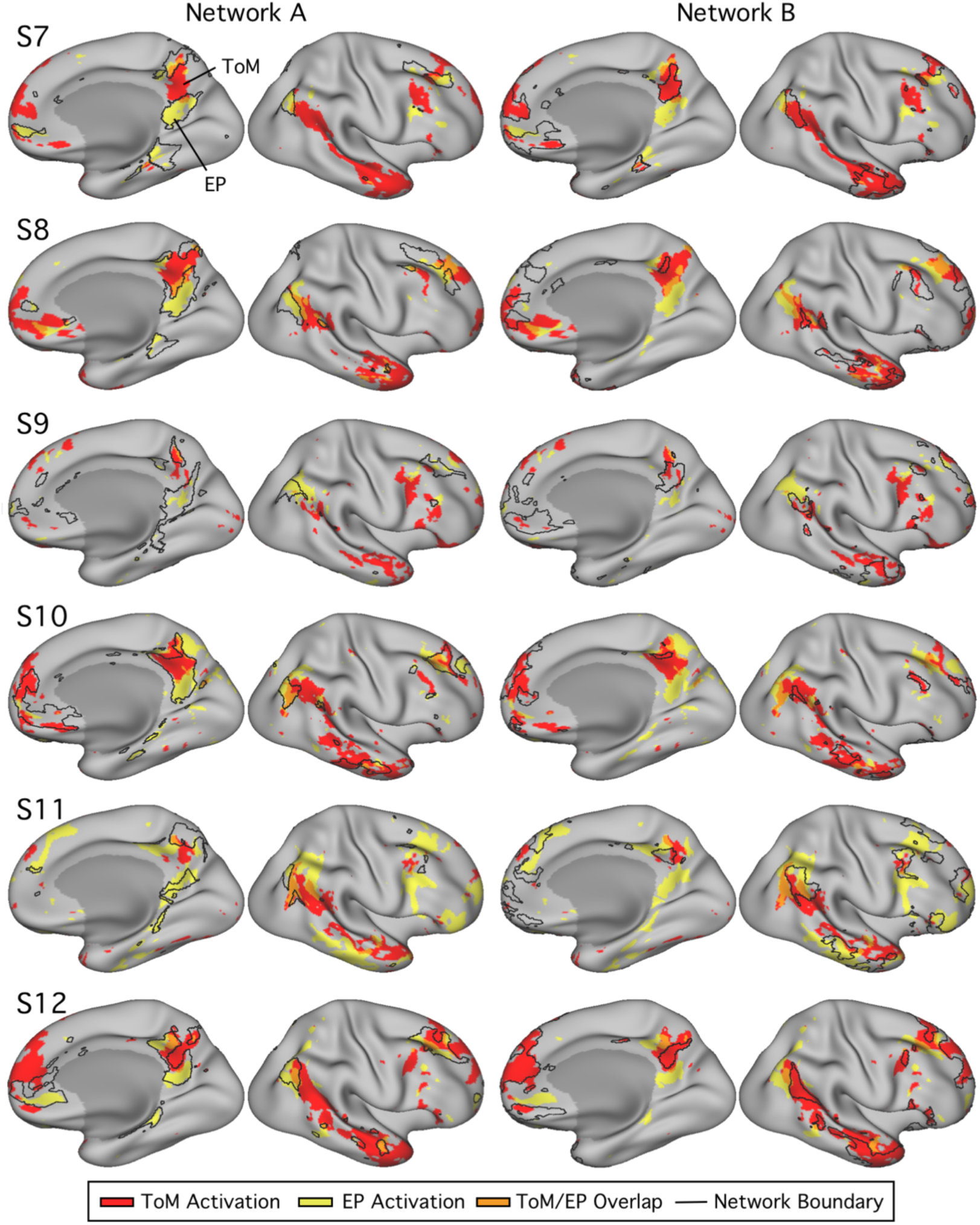
Networks A and B replicate differential recruitment by Episodic Projection and ToM tasks in the right hemisphere in additional individuals. Using the same procedures as displayed in Fig. 12, the right hemisphere is displayed for each of the 6 subjects from Exp. 2. Each row shows the lateral (right) and medial (left) views of the right hemisphere for all subjects, with Network A boundaries (left columns) or Network B boundaries (right columns).

These results replicate a functional dissociation of Networks A and B across distributed cortical zones, with Network A preferentially recruited for contrasts requiring Episodic Projection and Network B for contrasts requiring ToM.

### Functional Dissociation of Networks A and B Replicates Across Task Contrasts

Task-specific analyses again provided evidence for a functional double dissociation (Fig. 14). Within Network A, 5 of 6 subjects showed greater responses in each of the Episodic Projection task contrasts than either contrast of the ToM tasks (Cohen’s *d* range = 0.37-1.68 for pairwise comparisons; most *d* > 0.90). The anomalous individual’s response pattern (S10; see Fig. 10) was the same for both task contrasts within each domain, consistent with a subject – rather than a task – effect.

**Figure 14.**
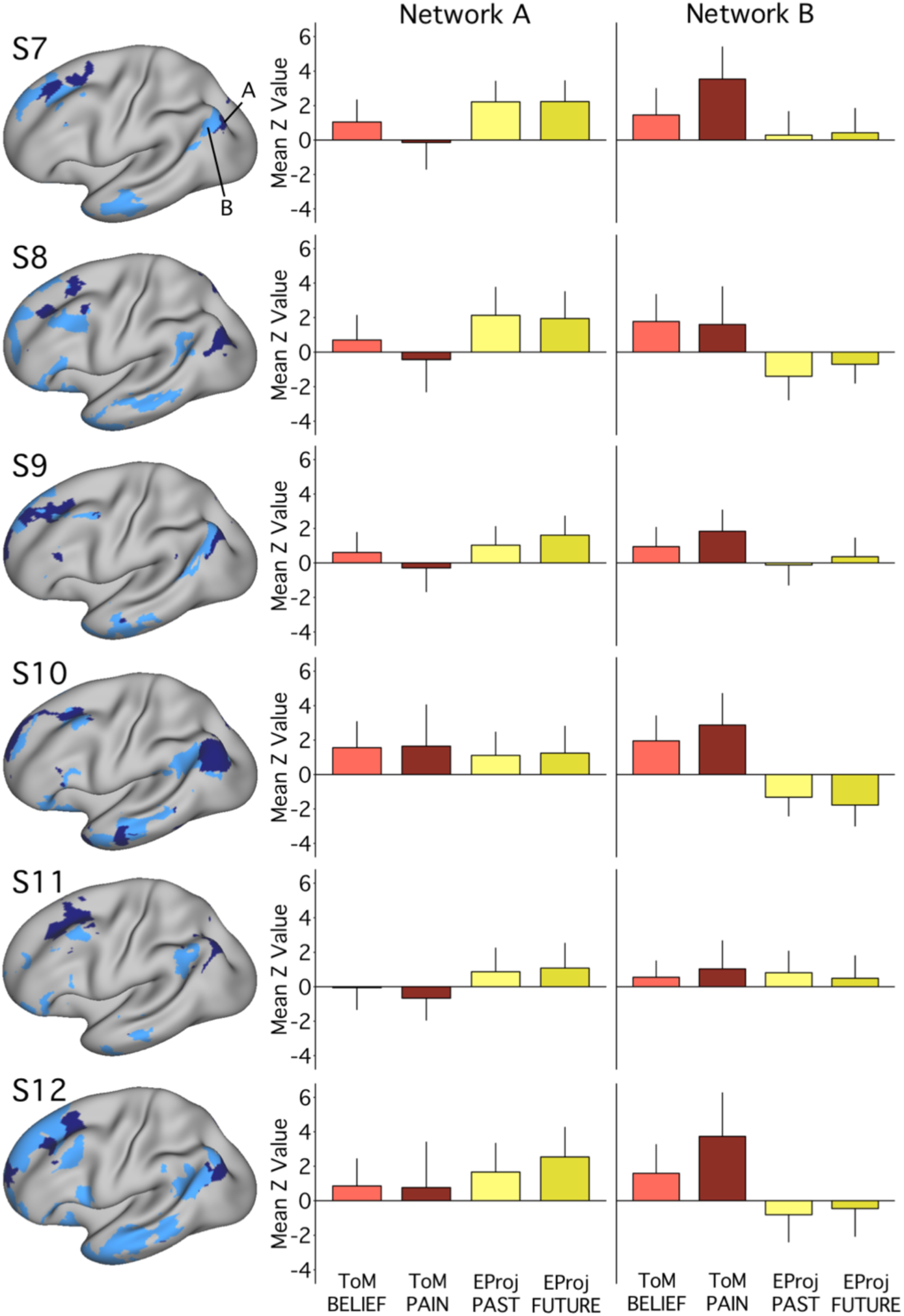
Functional dissociation of Networks A and B replicate across task contrasts in Exp. 2. The left column again shows the spatial distributions of Network A (navy) and Network B (light blue) for each individual in Exp. 2. The bar graphs plot the functional responses (mean *z*-values and standard deviations across runs) within each network for each specific task contrast. Replicating findings from Exp. 1, tasks within a domain exhibit comparable differences in network recruitment, and the Other Pain contrast shows the strongest recruitment of Network B in multiple individuals. For Network A, functional response increases for both Episodic Projection contrasts over both ToM contrasts are evident in 5 out of 6 subjects (Cohen’s *d* range = 0.37-1.68 for pairwise comparisons, most *d* > 0.90). For Network B, increases for both ToM contrasts over both Episodic Projection contrasts are evident in 5 out of 6 subjects (*d* range = 0.52-2.94, most *d* > 1.50).

Within Network B, as in Exp. 1, there was a systematic difference in the ToM domain, with a greater response for the ToM OTHER PAIN contrast relative to the FALSE BELIEF contrast in every subject except S8 (Cohen’s *d* range = 0.36-1.20). Yet, despite this difference, the ToM mean responses for both task contrasts were greater than the responses in either of the Episodic Projection contrasts for all subjects but S11 (*d* range = 0.52-2.94 for pairwise comparisons; most *d* > 1.50).

Thus, in all relevant ways, Exp. 2 replicated Exp. 1. In the few instances where individual subjects deviated, the patterns were consistent across specific task contrasts – the individual as a whole appeared atypical.

Motivated to examine additional facets of network recruitment, we next conducted an exploratory analysis of self-referential task contrasts that were not possible in Exp 1.

### Exploratory Analysis of Episodic Projection Contrasts Varying Time Orientation

We conducted an exploratory analysis of the three new contrasts from Exp. 2’s Episodic Projection paradigm that differed in temporal orientation: PAST SELF versus PAST NON-SELF, PRESENT SELF versus PRESENT NON-SELF and FUTURE SELF versus FUTURE NON-SELF. Since all three contrasts include SELF versus NON-SELF components, each contrast isolates a distinct temporal context. Admittedly any such complex contrast will inadvertently change other task features and is certainly imperfect. Nonetheless, analyzing the three separate contrasts allows for tests of generality, and where differences are found, seeds observations for future investigation. The PAST and FUTURE contrasts, which required episodic construction of previously experienced or imagined scenes, were predicted to preferentially recruit Network A (e.g., Andrews-Hanna et al. 2010), while all three contrasts, featuring self-referential component processes, were predicted to recruit Network B. The results are illustrated in Fig. 15.

**Figure 15.**
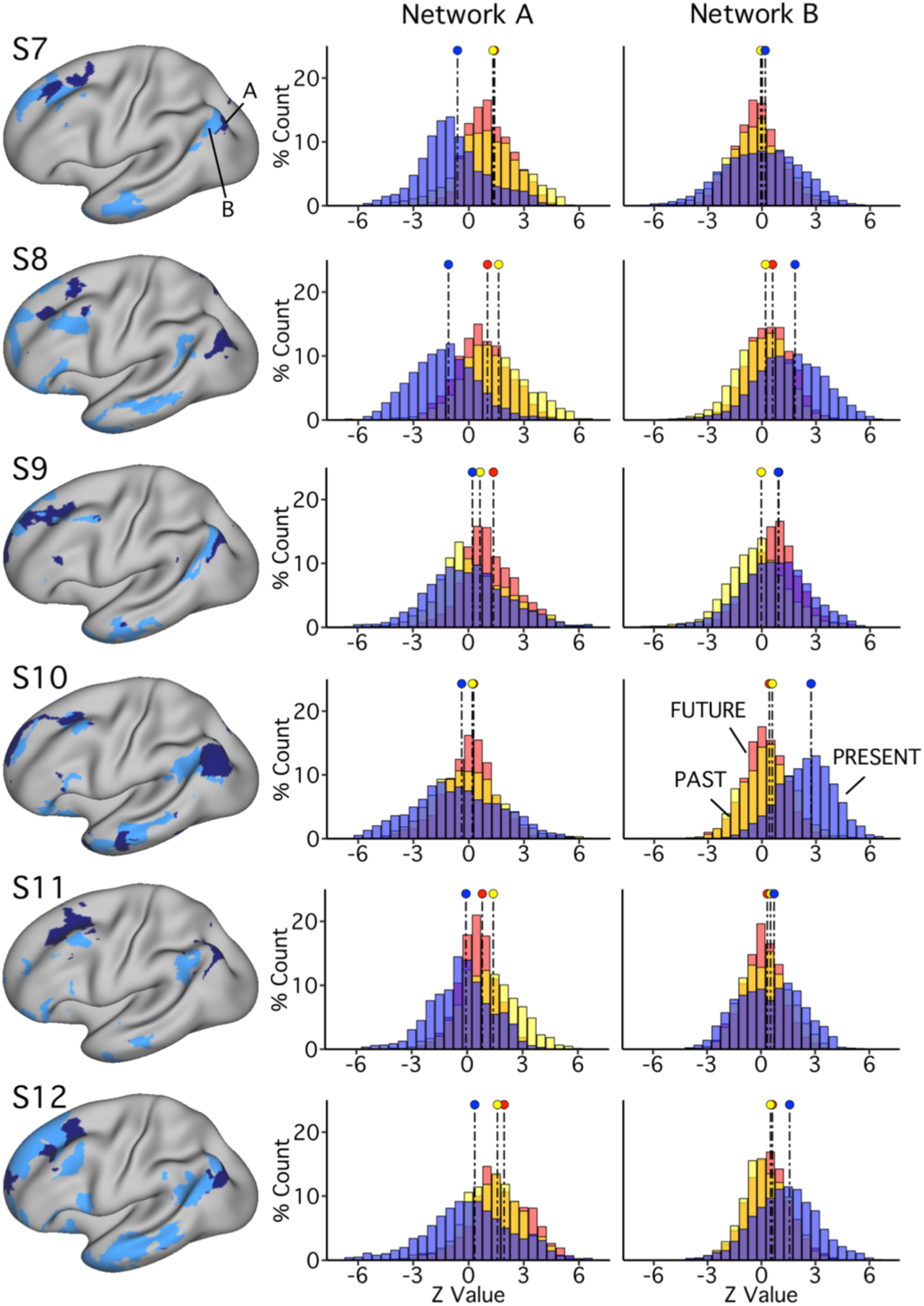
Exploratory analysis of network recruitment for self-referential tasks across temporal orientations. Results from the time-matched contrasts in the Episodic Projection task are displayed. The distributions in the right columns plot the functional responses within each network for these contrasts (PAST SELF versus PAST NON-SELF in yellow, PRESENT SELF versus PRESENT NON-SELF in blue, and FUTURE SELF versus FUTURE NON-SELF in red). For Network A, all subjects show greater responses in both the PAST and FUTURE conditions relative to the PRESENT (Cohen’s *d* range = 0.20-1.47; most *d* > 0.60). For Network B, the PRESENT condition shows a shifted response relative to the PAST and FUTURE conditions in almost all instances (with the one exception of the FUTURE condition in S9; *d* range = 0.10-1.67, most *d* > 0.50).

As predicted, for Network A, all 6 subjects exhibited greater recruitment by both the PAST and FUTURE contrasts over the PRESENT contrast, with 4 individuals (S7, S8, S11, S12) showing differences of medium to large effect (PAST+FUTURE > PRESENT: Cohen’s *d* range = 0.60-1.47 for pairwise comparisons). 3 of these subjects (S7, S8, and S12) were notable not only for how large the difference was between the temporally directed conditions (PAST and FUTURE) in comparison to the PRESENT contrast but also for how similar the two temporal conditions were to each other (Cohen’s *d* range = 0.05–0.36). Thus, the PAST and FUTURE contrasts elicit responses in Network A that are strong and largely similar.

For Network B, we did *not* observe the predicted, positive shift across the three self-referential contrasts. Rather, in almost all individuals (all but S9), the PRESENT contrast was greater than both the PAST and FUTURE contrasts, with 3 of the individuals (S8, S10, S12) showing a sizable effect (*d*=0.62-1.67). This result is consistent enough to warrant attention in future studies, but is not consistent with the prediction that self-referential episodic decisions would involve strong responses in both Networks A and B. A notable and unexpected finding is that the double-dissociation observed for the Episodic Projection contrasts appears to extend to PAST and FUTURE contrasts even when self-referential processing is not controlled.

A final observation for future consideration is quite subtle. In instances when an individual did not show contrast-related differences between conditions, little recruitment by *any* contrast was apparent, with the distributions all centered near zero (e.g., see S10 Network A and S7, S11 Network B). That is, there were no instances in which all three contrasts produced a clear positive response in either Network A or Network B.

## Discussion

Parallel networks at the transmodal apex of association cortex^1^, within the bounds of the canonical DN, can be defined within individual subjects and are preferentially recruited by distinct task domains. The PHC-linked Network A is preferentially recruited for task contrasts isolating Episodic Projection, including remembering the past and imagining the future, and the TPJ-linked Network B is preferentially recruited for task contrasts isolating ToM. The double dissociation between these two adjacent networks has practical implications for efforts to map and understand higher-order aspects of cognition. The spatial interdigitation of functionally specialized networks throughout distributed zones of cortex raises intriguing questions about their origins, including how such specialization arose through evolution and emerges during development.

### Parallel Distributed Networks with Adjacent Regions Across the Cortex Are Functionally Specialized

Overlap in group-analyzed task responses across a broad range of domains has driven ideas about the function and organization of the DN. Convergence has been highlighted for tasks targeting mnemonic through social functions (Buckner and Carroll 2007; Spreng et al. 2009; see also Mars et al. 2012). However, there have also been challenges to the idea of convergence. In an early critical study, Rosenbaum et al. (2007) demonstrated that performance on ToM tasks, as typically administered in neuroimaging studies, is preserved in patients with MTL damage, suggesting dissociation between episodic memory and ToM. Group studies that manipulate task domains within subjects note regional differences in task response patterns, as well, with a recurring distinction between activation in TPJ during ToM / mentalizing tasks in a zone rostral to the IPL region activated during episodic remembering (Andrews-Hanna et al. 2014a; DuPre et al. 2016; Spreng and Grady 2010). A particularly revealing analysis of this distinction is reported by Andrews-Hanna, Saxe, and Yarkoni (2014a). Using spatial meta-analysis that aggregated data from 79 task contrasts targeting mentalizing and 95 targeting episodic retrieval, they observed robust activation of the IPL for both tasks, but with the mentalizing response rostral to the memory response (Andrews-Hanna et al. 2014a).

In the current work, across two independent experiments, preferential responses in distinct distributed networks were observed across the brains of individuals for both of the task domains tested (ToM and Episodic Projection). Our findings thus reinforce the notion that there exists functional specialization of, rather than convergence on, processing nodes or networks recruited for these task domains. While the dissociation in the TPJ and PHC could be hypothesized from prior group analyses of similar tasks, it is striking that Networks A and B showed dissociations across the brain, including anterior and posterior midline zones previously described as components of a DN core (Andrews-Hanna et al. 2010; Andrews-Hanna et al. 2014b; Christoff et al. 2016; Tamir et al 2016).

In general, there was considerably more separation than overlap between the two task domains within most individuals. Particularly compelling evidence for a fine level of separation comes from visualization of the preferential task responses against the Network A and B boundary estimates in the clearest data sets (e.g., S2 in Fig. 6; S12 in Fig. 12). In these instances, the spatial separation of the task-preferential responses is distinct in both the anterior and posterior midline, consistent with separation of the two networks observed previously (Braga and Buckner 2017; Braga et al. 2019).

The distinct task activation patterns along the posterior midline are consistent with results from two recent studies that found similar differential activity within subjects (tasked with making judgments about either people or scenes; e.g., see Fig. 1 in Peer et al. 2015; see also Silson et al. 2019). The current results support that such task activation differences within canonical DN regions are well-predicted by the hypothesis of separation between two distributed networks, labeled Networks A and B, and are not limited to specific cortical zones. These findings encourage reconsideration of regions previously considered part of a “core” (or “hub”) interacting with multiple subnetworks. Rather, the present findings suggest that distributed, functionally-specialized networks may remain spatially separated across much (or even the entirety) of the cerebral mantle.

### Relationship Between Specialized Networks and Possible Developmental Origins

Dissociation between juxtaposed networks that share a similar organizational motif raises questions both about whether there are functional similarities or dependencies between the separate networks and about how such an organization might arise. While the present data are insufficient to offer strong conclusions, several possibilities and constraints can be discussed.

An interesting feature of the network organization revealed here is that it supports a parallel division of the canonical DN, rather than suggesting an orthogonal organization. This is a non-trivial point. Prior group analyses, due to spatial blurring, suggested an organization that blurred across the networks; higher spatial resolution now reveals a division of the canonical, group-averaged DN into two distinct networks. One possibility is that the two networks are domain-specialized but form a broader class of networks (Buckner and DiNicola In Press). Much like the visual system possesses specializations across multiple extrastriate areas and patches that process stimuli with distinct visual properties and experiential histories, higher-order association cortex might specialize similarly. Within this possibility, the two networks may possess broadly similar processing modes, perhaps aligned with internal mentation as opposed to external forms of perception, but also specialization for separate domains, here captured by our contrasts targeting ToM as distinct from Episodic Projection. This raises one of the most intriguing questions: how do the separate networks form?

Developmental specialization of higher order visual regions provides potential insight. Early during development, just after birth, extrastriate visual regions in the monkey do not possess fine specialization for stimulus domains (e.g., faces versus places) but do possess a broad retinotopic organization, by which central to peripheral portions of the retina are mapped to adjacent zones of cortex (Arcaro and Livingstone 2017a). As development progresses, face response patches form preferentially in extrastriate zones that are aligned to the central, as opposed to the peripheral, retinotopic maps. The early retinotopic proto-organization may, thus, serve as a scaffold onto which experience-based sculpting of the visual system is refined (Arcaro and Livingstone 2017a; see also Arcaro and Livingstone 2017b).

Refinement of association networks may similarly derive hierarchically from an early proto-organization that progresses from a broadly distributed DN-like network, in the earliest developmental stages, to multiple parallel networks at later stages, perhaps through a competitive developmental process of fractionation and specialization (Buckner and DiNicola In Press). Such a possibility would explain both the similarity of the two networks’ organization as well as their juxtaposition across the cortex. A key difference between the two networks, in most individuals, is that Network A is strongly coupled to posterior PHC, while Network B is not. This difference, perhaps reflecting some form of early limbic projection gradient onto cortex, may contribute to the initiation of specialization much as central-to-peripheral inputs from the retina scaffold specialization of extrastriate visual cortex.

Limitations to interpretation should also be noted. Specifically, our results should not be interpreted to mean ToM tasks exclusively utilize Network B or that tasks involving remembering and prospection exclusively utilize Network A. Here, task *contrasts* were selected based on prior literature because they preferentially activated components of one or the other of the two networks and did so through subtraction. This highlights the need for caution against over-interpreting the results as suggesting that the networks are recruited selectively in an absolute sense. What we illustrated in our primary analyses, by selecting task contrasts that isolated component processes of ToM as distinct from processes of Episodic Projection, is a functional dissociation between the two networks that is present across the distributed network regions. Less-constrained task contrasts are likely to produce less-differentiated results and may be expected to call upon both Networks A and B, although preliminary analyses of additional task contrasts in Exp. 2 provided *more* evidence for functional specialization, not less (Fig. 15).

### Individual Differences

Within our primary analysis of network recruitment by task domain, some subjects revealed clearer dissociations than others. It is unclear in the present results whether there are biologically meaningful differences in the patterns across subjects or we are simply exploring patterns at the resolution edge of our methods. In Braga and Buckner (2017), the distinction between fine scale networks was more robust in two of four individuals, and in Braga et al. (2019), as well, a subset of individuals provided the most robust data, thus showing similar variability to the present findings. Future work and larger numbers of subjects will be required to understand sources of individual differences.

## Conclusions

The present work shows that recently identified Networks A and B, within the bounds of the canonical DN, can be functionally dissociated by task contrasts from distinct domains of internal mentation. Network A is preferentially recruited by task contrasts isolating episodic projection, including remembering the past and imagining the future. Network B is preferentially recruited by task contrasts isolating theory of mind. Questions remain as to the full extent of the functional specialization of these networks, but the present results encourage a re-evaluation of how Networks A and B are organized to support task processing demands.

## Acknowledgements

We thank T. O’Keefe, H. Hoke, R. Mair and S. McMains for assisting in data acquisition and preprocessing optimization. E. Phlegar, M. Shermohammed, B. Braams, C. Vidal Bustamante, and T. Moran also assisted in data acquisition. A. Youssoufian, H. Becker, K. Miclau and A. Song assisted in data acquisition and stimuli generation. We thank the Harvard Center for Brain Science neuroimaging core and FAS Division of Research Computing. The multi-band EPI sequence was generously provided by the Center for Magnetic Resonance Research (CMRR) at the University of Minnesota. Supplemental task stimuli were provided by R. Saxe and J. Andrews-Hanna, and R. Saxe provided valuable discussion during the project.

## Footnotes

1 Analyses of the hierarchical organization of networks in both humans and monkeys have converged on a macroscale gradient that progresses outwards from primary sensory-motor cortex through to higher-order association regions (Buckner and DiNicola In Press; Buckner and Margulies 2019; Huntenburg, Basin, and Margulies 2018; Margulies et al. 2016; see also Mesulam 1998). The historically-defined default network is spatially positioned at the apex of the gradient, encompassing distributed transmodal association regions including frontopolar and rostral regions of temporal association cortex.

